# Discovery of two archaeal GDGT lipid modifying enzymes reveals diverse microbes capable of H-GDGT biosynthesis and modification

**DOI:** 10.1101/2023.10.20.563219

**Authors:** Andy A. Garcia, Grayson L. Chadwick, Paula V. Welander

## Abstract

Archaea produce unique membrane-spanning lipids, termed glycerol dialkyl glycerol tetraethers (GDGTs), which are thought to aid in adaptive responses to various environmental challenges. GDGTs can be modified in a variety of ways, including cyclization, bridging or cross-linking, methylation, hydroxylation, and desaturation, to give rise to a plethora of structurally distinct GDGT lipids with different properties. Here we report the discovery of a pair of radical SAM enzymes responsible for two of these modifications - an H-GDGT bridge synthase (Hbs), responsible for cross-linking the two hydrocarbon tails of a GDGT to produce H-GDGTs and an H-GDGT methylase (Hgm), responsible for the subsequent methylation of H-GDGTs. Heterologous expression of Hbs proteins from various archaea in *Thermococcus kodakarensis* results in the production of H-GDGTs in two isomeric forms. Further, co-expression of Hbs and Hgm results in the production of mono- and di-methylated H-GDGTs and minor amounts of tri-methylated H-GDGTs while expression of Hgm alone results in minor production of mono- and di- methylated GDGTs. Phylogenetic analyses reveal the presence of Hbs homologs in diverse archaeal genomes spanning all four archaeal superphyla. We also find Hbs homologs in bacterial genomes that have the genetic potential to synthesize fatty acid-based membrane-spanning lipids such as brGDGTs. We subsequently demonstrate H-GDGT production in three Hbs-encoding archaea, identifying an increase in H-GDGTs in response to elevated temperature in members of the genus *Archaeoglobus* and observing the production of highly cyclized H-GDGTs with up to 6 rings in the Thermoproteales archaeon *Vulcanisaeta distributa.* Such highly cyclized H-GDGTs are the precursors of ARN acids, a class of tetraprotic naphthenic acids that cause destructive mineral deposition during crude oil processing. Co-occurrence of the H-GDGT synthase with the previously identified GDGT ring synthases in archaeal genomes allowed identification of multiple archaeal phyla with the genetic potential to produce highly cyclized H-GDGTs, with particularly interesting candidates in the class Thermoplasmata from oil rich environments.

## Introduction

Archaea are a unique group of microbes that differ from bacteria and eukaryotes in a multitude of ways with one of the most prominent distinctions being the composition of their membrane lipids. Archaea possess structurally distinct bilayer membrane lipids known as archaeols (or diethers) which are made from glycerol-1-phosphate ether bonded to a pair of C_20_ isoprenoid-based (phytane) hydrocarbon tails (1). In contrast, bacterial and eukaryotic membrane lipids are composed of glycerol-3-phosphate ester bonded to fatty acid-derived tails. Many archaea also fuse their bilayer lipids into a monolayer of membrane spanning lipids, known as glycerol dialkyl glycerol tetraethers (GDGTs) (2). Although less widespread, bacteria also fuse their phospholipid bilayers to form analogous monolayer lipids, a subclass of which are termed branched GDGTs (brGDGTs) (3). GDGT lipids can be further modified through cyclization (formation of cyclopentane or cyclohexane rings), cross-linking between the two alkyl chains, methylation, hydroxylation, and desaturation (2, 4). These GDGT modifications have been shown in culture experiments to aid in survival under extreme environmental conditions such as high temperatures, acidic pH, or low nutrient availability (5–9). Additionally, the recalcitrant nature of GDGTs results in their preservation in the rock record over hundred-million-year timescales where they can function as geological proxies used to reconstruct paleoenvironmental conditions (10). Modified GDGTs, such as cyclized GDGTs, have proven to be some of the most useful paleotemperature proxies contributing to our understanding of changes in atmospheric and sea surface temperatures over geological timescales (11–14).

One of the more unusual modifications observed in GDGT biosynthesis is the formation of H-GDGTs, also referred to as glycerol monoalkyl glycerol tetraethers (GMGTs) (15). These lipids are characterized by a covalent bond between the two alkyl chains of a GDGT molecule (Fig. 1 and Fig. S1). The exact location of this bond is not known with certainty but is suggested to occur between the C-20 and C-20” methyl carbons of a GDGT (Fig. S1) (16, 17). H-GDGTs were first discovered in two thermophilic methanogens, *Methanothermus sociabilis* and *Methanothermus fervidus,* and have since been found in a variety of other thermophilic Euryarchaeota within the phyla Methanococci, Methanopyri, Thermococcales, and Thermoplasmatota as well as one TACK archaeon within the Thermoprotei (15, 18–24). H-GDGTs can possess additional modifications including cyclopentane rings as seen in *Aciduliprofundum boonei* and *Ignisphaera aggregans*, amongst others, as well as methylations which are only observed in a few thermophilic methanogens including *Methanopyrus kandleri* and *Methanothermobacter thermautotrophicus* (18, 19, 25, 26). Structurally diverse H-GDGTs have also been detected directly in a variety of environments, including hot springs, hydrothermal vents, lake sediments, and marine sediments (27–30) These environmental H-GDGTs include ring containing H-GDGTs, H-GDGTs with 1-3 methylations, and bacterial derived branched H-GDGTs (brH-GDGTs) (25, 29, 31). Furthermore, archaeal H-GDGTs occur in two isomeric forms (termed H-GDGT-0a and H-GDGT-0b in this work), but the structural basis of these isomers is currently unresolved (32).

**Figure 1.**
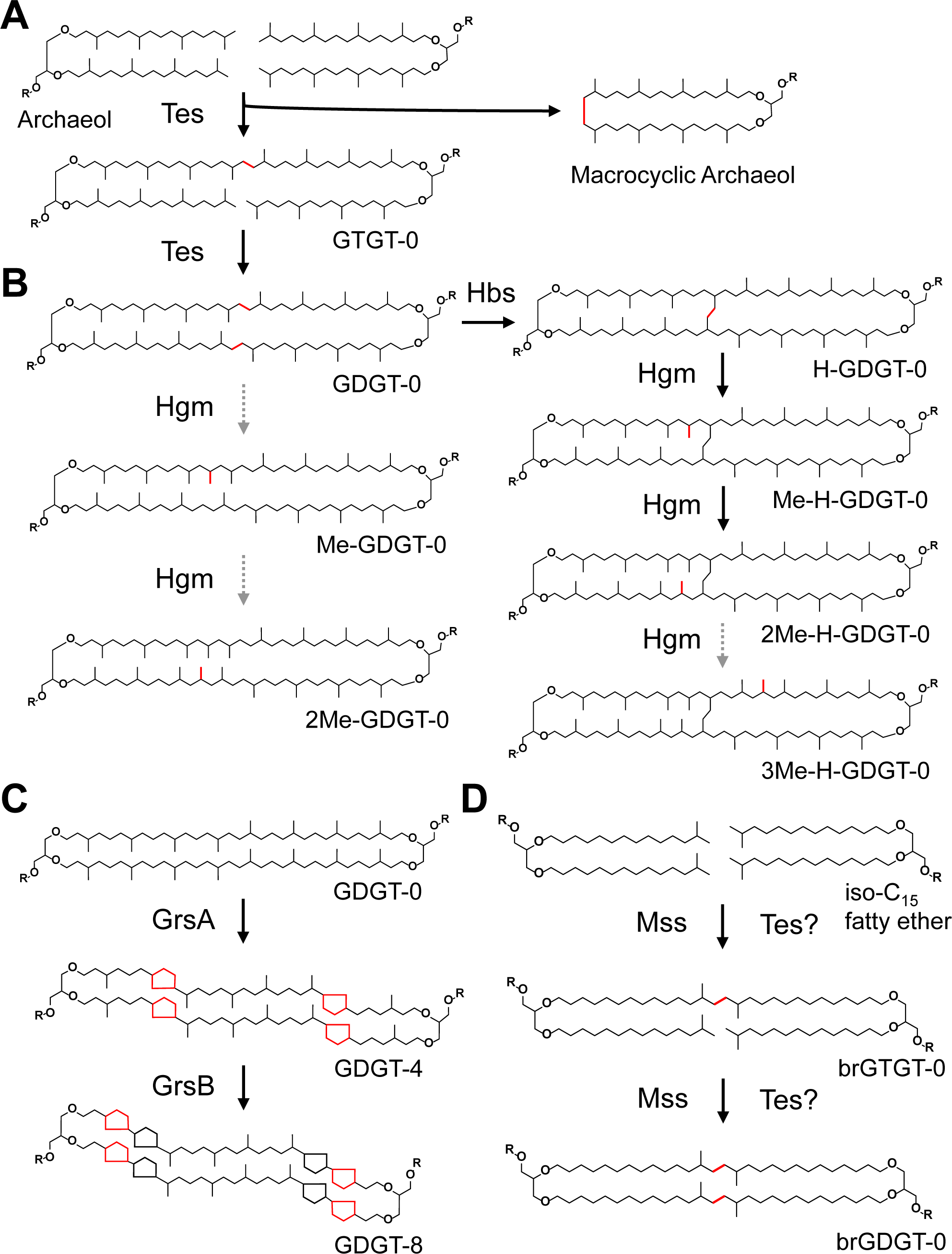
Radical SAM proteins are involved in archaeal and bacterial GDGT biosynthesis. A) Two molecules of archaeol are dimerized to form a GDGT molecule by Tes which also produces the intermediate GTGT-0 and the side product macrocyclic archaeol. B) The biphytanyl chains within a GDGT are bridged/cross-linked by the newly discovered Hbs protein to form H-GDGTs which are then methylated by Hgm to form mono-, di-, and tri-methylated H-GDGTs. Hgm can also methylate unbridged GDGTs to yield mono- and di-methylated GDGTs. The positions of the bridge and methylations are tentatively assigned based upon previous NMR studies of ARN acids (REF) but have not yet been confirmed in intact lipids. C) Cyclized GDGTs are synthesized by the related GrsA and GrsB enzymes which introduce up to four cyclopentane rings at the C-7 and C-3 positions of a GDGT molecule, respectively. D) Bacterial branched GDGTs are synthesized by Mss and potentially also Tes via the dimerization of bacterial bilayer lipids based on iso-C_15_ fatty acids. GDGT: glycerol dialkyl glycerol tetraether; GTGT: glycerol trialkyl glycerol tetraether; Tes: tetraether synthase; Hbs: H-GDGT bridge synthase; Hgm: H-GDGT methylase; Grs: GDGT ring synthase; Mss: membrane-spanning lipid synthase.

From a physiological perspective, the benefit of generating a bridged GDGT remains unclear. The observation that H-GDGTs are predominantly produced by thermophilic archaea and are most abundant in extreme environments (30), has led to the proposal that H-GDGTs play an important role in membrane adaptation to harsh and fluctuating environmental conditions. Environmental studies have shown a positive correlation between H-GDGT abundances and increasing temperatures in marine sediment and peat (30, 33). In culture studies, environmental stress factors other than increased temperature have been shown to have an impact on H-GDGT production as well. One study demonstrated that *Pyrococcus furiosus*, a hyperthermophilic marine archaeon, increased the percentage of H-GDGTs as a response to acidic pH, high salinity, or decreasing temperature, which can be a significant stress factor in a hyperthermophile (21). However, the physiological advantage imparted by H-GDGT formation under these conditions is not known. From a geochemical perspective, the diagenetic products of H-GDGTs have been identified in hundred-million-year-old sediments dating back to the Jurassic period (34) and are also thought to be the precursor to ARN acids – tetraacid compounds found within crude oils that cause destructive precipitation of calcium naphthenate minerals (16, 17, 35). However, the use of H-GDGTs as paleoproxies and their contribution to petroleum geochemistry remain unresolved as more insights are needed on the potential biotic sources of H-GDGTs, their preservation potential, and in the underlying molecular mechanisms linking environmental adaptation to membrane modifications in H-GDGT producers.

One current obstacle to comprehending the physiological and geochemical significance of H-GDGTs is our lack of information on how H-GDGTs are synthesized and modified in the cell. While many diverse structures of GDGTs are known (2), the enzymes responsible for their synthesis and modifications have only recently been discovered. The formation of archaeal GDGTs was shown to be carried out by the enzyme tetraether synthase (Tes), a radical S-adenosylmethionine (SAM) protein that links two archaeol molecules to generate a GDGT (Fig. 1) (36, 37). Similarly, the synthesis of bacterial monolayer lipids is carried out by another radical SAM enzyme, membrane spanning lipid synthase (Mss) and potentially also Tes homologs (Fig. 1) (37, 38). Further, the B12-binding radical SAM proteins, GDGT ring synthases (GrsA and GrsB), were shown to be responsible for the formation of cyclopentane rings at different locations within the biphytane core of GDGTs (Fig. 1) (39). In some thermoacidophilic archaea, GDGT hexose head groups are converted to the five-membered ring structure known as calditol by the calditol synthase (Cds), another radical SAM enzyme (40) . With the knowledge that four enzymes (Tes, Mss, Grs, and Cds) involved in GDGT biosynthesis belong to the radical SAM superfamily, we hypothesized that radical SAM enzymes would also be required for the formation and methylation of H-GDGTs.

In this study, we confirm this hypothesis through the identification of an H-GDGT bridge synthase (Hbs) belonging to the radical SAM SPASM/Twitch subfamily that is responsible for forming the covalent bond between the two biphytanyl chains of a GDGT molecule. Additionally, we confirm that an uncharacterized B12-binding radical SAM protein that often clusters with the H-GDGT bridge synthase gene in a variety of archaeal genomes encodes an H-GDGT methylase (Hgm). Through bioinformatic analyses, we identify Hbs homologs in the genomes of diverse archaea spanning the Asgard, DPANN, Euryarchaeota, and TACK superphyla and in bacteria possessing Mss and Tes homologs, suggesting that Hbs homologs may also be involved in the biosynthesis of bridged membrane spanning lipids in bacteria, such as brH-GDGTs. We also look to cultured archaea to study native H-GDGT production, observing substantial increases in the relative abundance of H-GDGTs in response to increased temperature in the genus *Archaeoglobus* and identifying *Vulcanisaeta distributa* as the first cultured archaeon capable of producing highly cyclized H-GDGTs with 5 and 6 rings. Taken together, our study demonstrates the utility of identifying key GDGT biosynthesis proteins to characterize the unique biochemistry and physiological significance of lipid modifications in archaea.

## Results and Discussion

### Identification of a H-GDGT bridge synthase protein

To identify proteins responsible for H-GDGT biosynthesis, we searched for radical SAM enzymes that are shared by three H-GDGT producers, *Thermococcus celer*, *Thermococcus guaymasensis*, and *Pyrococcus furiosus,* and are absent in a closely related archaeon, *Thermococcus kodakarensis,* that does not produce H-GDGTs. Our search revealed a radical SAM protein that was shared among the three H-GDGT producers that had significant similarity (e-value = 7e-79, 55% similarity, 36% identity) to the biochemically characterized *Methanosarcina barkerii* heme decarboxylase protein, AhbD (41). However, closer inspection of the AhbD homologs shared by the three H-GDGT producers revealed that they have an approximately 100 amino acid N-terminal extension, annotated as COG4001, which is absent in the *M. barkerii* AhbD protein (Fig. 2A). Furthermore, AlphaFold models (42, 43) predict that this extension forms a distinct domain and possesses a new binding pocket that is lined with hydrophobic residues necessary for binding lipids and is directly connected to the active site via a tunnel(s) (Fig. 2B and Fig S2-S4). Thus, we hypothesized that the AhbD homologs identified in the three H-GDGT producers were H-GDGT bridge synthases rather than heme decarboxylases.

**Figure 2.**
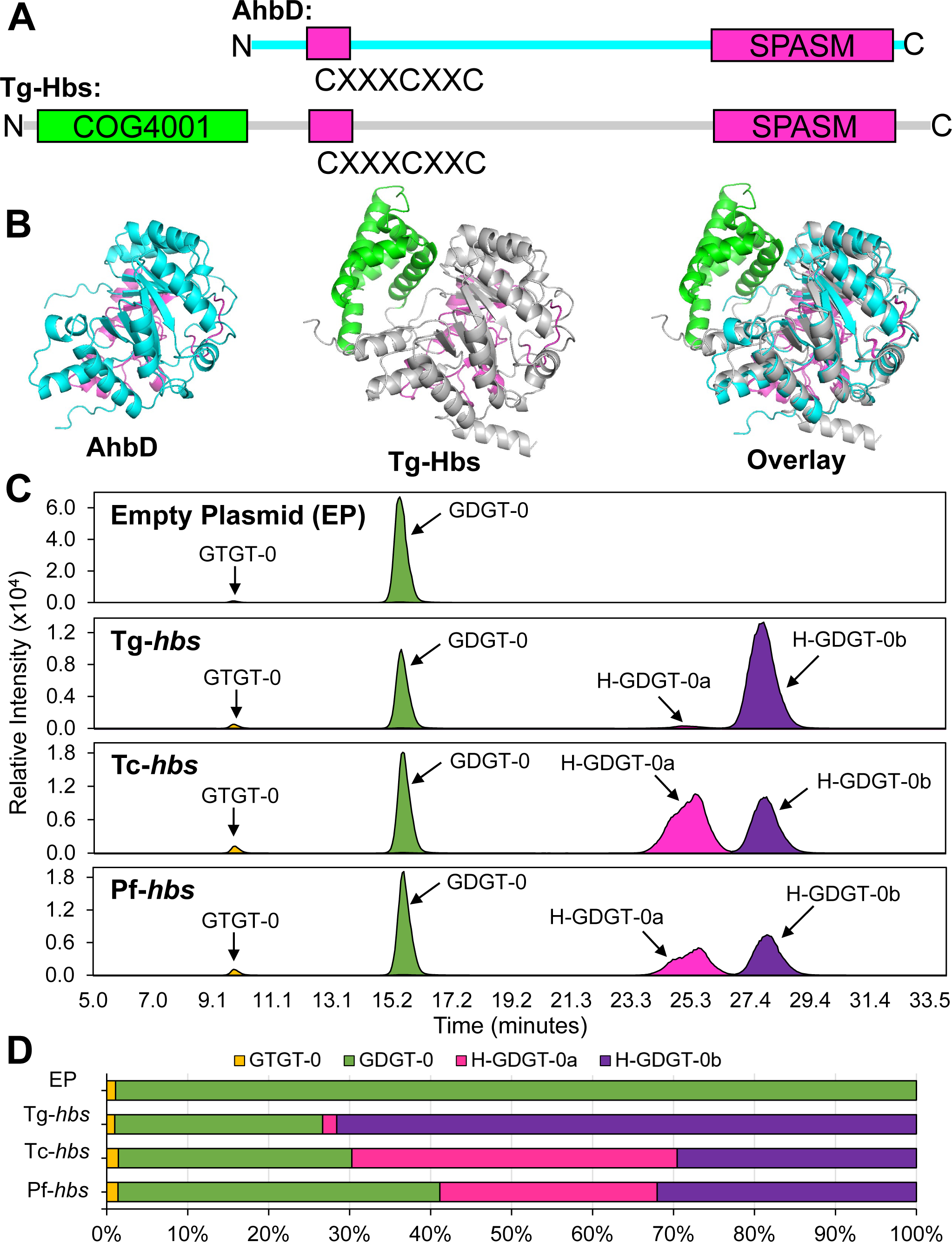
**The Hbs synthesizes H-GDGTs in two isomeric forms**. A) Hbs and the heme decarboxylase, AhbD, are related proteins which belong to the SPASM/Twitch subfamily of radical SAM enzymes. They possess the characteristic iron-sulfur cluster binding motif - CXXXCXXC - as well as 7-8 additionally conserved cysteine residues in their C-terminus (the SPASM domain) which are responsible for binding two auxiliary iron-sulfur clusters. Hbs additionally possess an ∼100 amino acid N-terminal extension annotated as COG4001, a sequence group with no known function, which is absent in AhbD. B) Alpha fold structure of Hbs predicts that the COG4001 sequence forms a new domain on the protein which shares no structural homology with AhbD. C) Merged extracted ion chromatograms (EICs) showing that the heterologous expression of Hbs proteins in *Thermococcus kodakarensis* results in robust H-GDGT production. The parental strain possessing an empty plasmid (EP) makes only GTGT-0 and GDGT-0. Strains expressing Tg-*hbs*, Tc-*hbs*, and Pf-*hbs* all produce H-GDGTs in two isomeric forms – H-GDGT-0a and H-GDGT-0b. D) Relative abundance of monolayer lipids in the control and heterologous expression strains showing that H-GDGTs are the most abundant monolayer lipids in strains expressing Hbs proteins, and that the two H-GDGTs isomers are present in largely different ratios depending on which Hbs protein is expressed with Tg-*hbs* predominately producing H-GDGT-0b and Tc-*hbs* and Pf-*hbs* producing near 1:1 mixtures of both isomers.

To determine if these candidate H-GDGT bridge synthases (Hbs) can bridge or cross-link GDGT alkyl chains to form H-GDGTs, we placed each candidate gene from *T. guaymasensis* (Tg-Hbs), *T. celer* (Tc-Hbs), and *P. furiosus* (Pf-Hbs) under the control of a strong promoter on the self-replicating plasmid pTS543 (44) and then introduced each plasmid into the heterologous host *T. kodakarensis*. We then acid hydrolyzed and extracted the core lipids from these cultures in triplicate and characterized the lipid extracts through liquid chromatography-mass spectrometry (LC-MS) in normal phase. The control parental *T. kodakarensis* strain (AL010) containing an empty plasmid produced GDGTs, a minor amount (∼1% of monolayer lipids) of glycerol trialkyl glycerol tetraethers (GTGTs), and no H-GDGTs (Fig. 2C and 2D). In contrast, heterologous expression of the three Hbs homologs in the parental *T. kodakarensis* strain resulted in robust H-GDGT production (Fig. 2C and 2D). H-GDGTs were the most abundant monolayer lipid in these strains, comprising 73%, 70%, and 59% of monolayer lipids in AL010 + Tg-*hbs*, AL010 + Tc-*hbs*, and AL010 + Pf-*hbs*, respectively. Each Hbs protein produced a mixture of the H-GDGT-0a and H-GDGT-0b isomers but in different proportions. Tg-Hbs produced 40 times more H-GDGT-0b than H-GDGT-0a while Tc-Hbs and Pf-Hbs both produced a near 1:1 mixture of each H-GDGT isomer (Fig 2D and Fig S5). Further, the ratio of the H-GDGT isomers produced by each Hbs protein was consistent and distinctive: 41.3±0.6 (Tg-Hbs), 0.7±0.01 (Tc-Hbs), and 1.2±0.01 (Pf-Hbs). While the structural difference between H-GDGT-0a and H-GDGT-0b is not known, it is possible that the two H-GDGT isomers differentially affect the biophysical properties of the membrane. We observed a significant increase (p < 0.05) in the bilayer to monolayer lipid ratio in the strains expressing Hbs proteins compared to the control strain and that this increase was significantly more for Tg-*hbs* than in Tc-*hbs* or Pf-*hbs*. (Fig. S5B). This suggests that *T. kodakarensis* may be adaptively responding to the unnatural presence of H-GDGTs in its membrane by decreasing monolayer lipid production and may be doing so to differing degrees based on the isomeric composition of the H-GDGTs present. Altogether, these data show that the extended AhbD homologs possessing COG4001 domains are H-GDGT bridge synthases that produce a mixture of two H-GDGT isomers.

### Identification of a H-GDGT methylase protein in *Thermococcus guaymasensis*

Inspection of the genomic neighborhood of the *hbs* gene in *T. guaymasensis* revealed that *hbs* occurs in a gene cluster with a GDGT ring synthase (*grs*) gene, and a gene encoding an uncharacterized B12-binding radical SAM protein (Fig 3A). This uncharacterized radical SAM protein was found to share similarity with the Moek5 methylase (e-value = 9e-63, 52% similarity, 31% identity), a protein involved in the biosynthesis of Moenomycin, an antibiotic that bears structural similarity to archaeal lipids (45). Additionally, the clustering of *hbs* genes with homologs of this putative methylase was found to occur in many divergent archaeal genomes, including in Nitrososphaerota, Thermoprotei, Thorarchaeota, and the three Euryarchaeota known to produce methylated H-GDGTs: *Methanopyrus kandleri, Methanothermobacter thermautotrophicus,* and *Methanothermobacter marburgensis* (Fig 3A). Taken together, we hypothesized that this gene encodes an H-GDGT methylase (Hgm).

**Figure 3.**
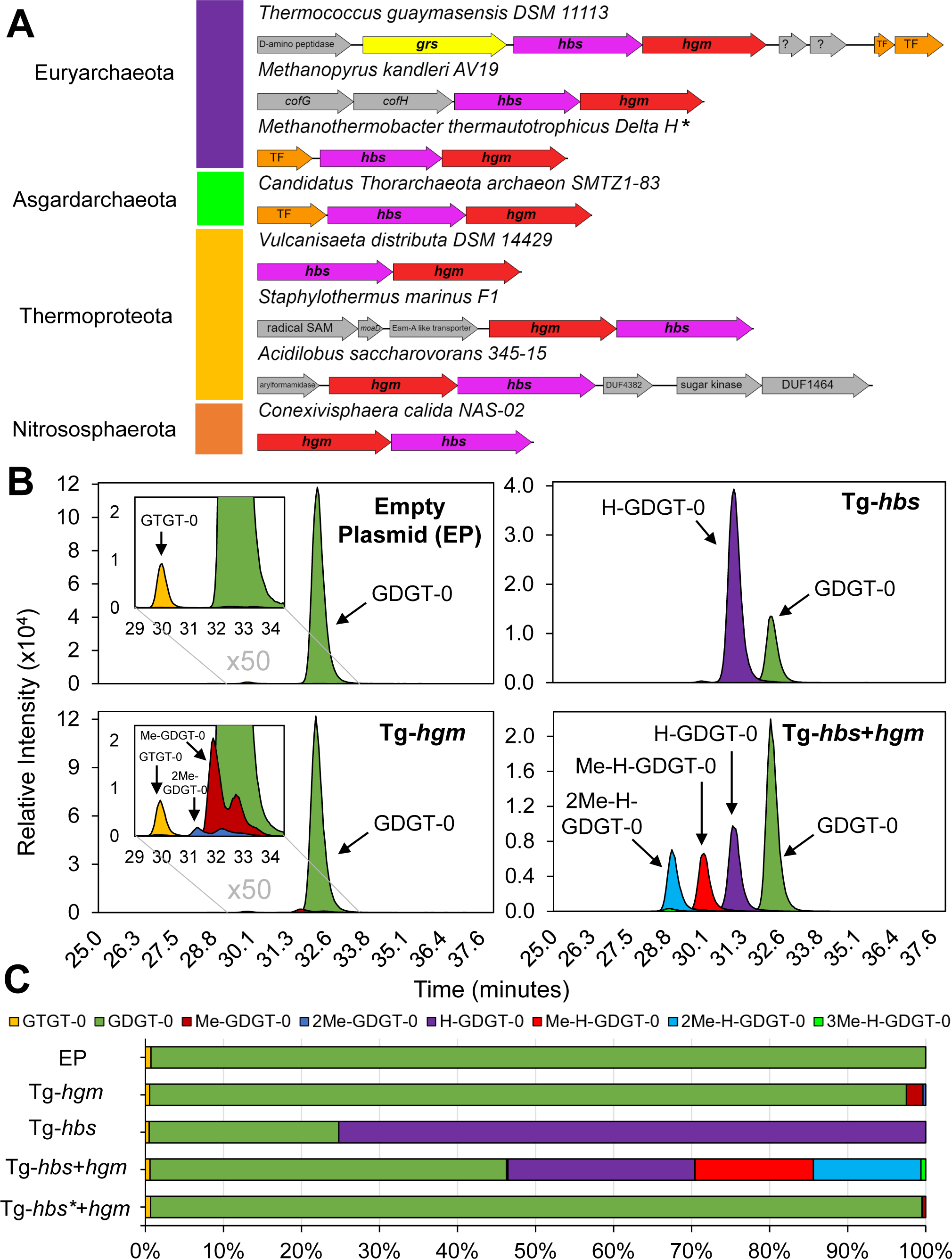
Hgm methylates H-GDGTs and, to a lesser extent, unbridged GDGTs. A) Hbs genes from diverse archaeal phyla occur in gene clusters with a putative H-GDGT methylase. *Methanopyrus kandleri* and *Methanothermobacter thermautrophicus* are known to produce methylated H-GDGTs. *The *M. thermautotrophicus hbs* gene cluster is the same as that in *M. marburgensis,* which also produces methylated H-GDGTs. B) Merged EICs showing that heterologous expression of Tg-*hgm* alone and with Tg-*hbs* in *T. kodakarensis* results in the production of methylated monolayer lipids. Expression of Hgm alone results in the minor production of mono- and di-methylated GDGTs. Co-expression of Hgm and Hbs results in the robust production of mono- and di-methylated H-GDGTs and trace tri-methylated H-GDGTs. C) Relative abundance of monolayer lipids in the control and heterologous expression strains, demonstrating that the majority of H-GDGTs are methylated in the Hgm+Hbs co-expression strain while only a minor amount of GDGTs are methylated in the control strain, in the Hgm alone, and in the catalytically inactivated Hbs (Hbs* + Hgm) co-expression strain.

To test if the *T. guaymasensis* methylase homolog was capable of methylating H-GDGTs, we co-expressed the putative Hgm from *T. guaymasensis* (Tg-Hgm) together with the H- GDGT biosynthesis protein Tg-Hbs. The expression cultures were acid hydrolyzed and core lipids were extracted in triplicate and analyzed by LC-MS in reverse phase to achieve better separation of methylated monolayer lipids. Co-expression of Tg-Hgm and Tg-Hbs resulted in the robust production of mono- and di-methylated H-GDGTs and a minor amount of tri-methylated H-GDGTs while the control strain expressing Tg-Hbs alone produced only non-methylated H-GDGTs (Fig. 3B and Fig. S6). Approximately 45% of H- GDGTs were unmethylated, 28% mono-methylated, 26% di-methylated, and 1% tri- methylated in the Tg-Hbs-Hgm co-expression strain (Fig 3C). The H-GDGT-0a and H- GDGT-0b isomers co-elute in reverse phase, but analysis of the Tg-Hbs-Hgm co- expression strain lipids with normal phase chromatography revealed mono-, di-, and tri- methylation on the H-GDGT-0b isomer and only mono-methylation on the much less abundant H-GDGT-0a isomer (Fig. S7). These methylations take place on the alkyl chain of the H-GDGT and not the glycerol backbone as confirmed by MS/MS spectra and the co-elution of the methylated and un-methylated H-GDGTs during normal phase chromatography (Fig. S8-S10). The exact position of the methylations within the alkyl chain was not determined definitively but are tentatively assigned at C-13 based on previous structural investigations on methylated GDGTs and methylated ARN acids (17, 26) Interestingly, methylation of un-bridged GDGTs occurred in trace amounts in the Tg-Hbs- Hgm co-expression strain. We measured 99.6% of the GDGTs as unmethylated while 0.3% were mono-methylated and 0.2% were di-methylated, demonstrating Hgm can utilize un-bridged GDGTs as a substrate. Furthermore, the expression of Tg-Hgm in the absence of Hbs and H-GDGTs resulted in more, but still minor, production of methylated GDGTs: 97.5% unmethylated GDGTs, 2.1% mono-methylated GDGTs, and 0.4% di- methylated GDGTs (Fig 3B and 3C). Previous studies have reported trace amounts of methylated GDGTs in wild type *T. kodakarensis* (46, 47). We also observed trace amounts of mono- and di-methylated GDGTs in the empty plasmid control, however, they were substantially less than in the Tg-Hgm expression strain (<0.1% mono-methylated GDGT and <0.1% di-methylated GDGT) and eluted differently than the methylated GDGTs seen during Hgm expression (Fig. S11 and S12). These data suggest that Hgm is an H-GDGT methylase that either prefers to methylate H-GDGTs over un-bridged GDGTs and/or requires an interaction with Hbs to function properly. If interaction with Hbs is required, we hypothesized that Hgm would methylate un-bridged GDGTs more robustly in the presence of such an interaction, especially if no competing H-GDGT substrate was present. To test this, we constructed a *T. kodakarensis* strain co-expressing Hgm and a catalytically inactivated Hbs where the radical SAM iron-sulfur cluster binding motif CXXXCXXC was mutated to AXXXAXXA. Analysis of the lipids in this strain (AL010 + Tg- Hbs*-Hgm) revealed only trace GDGT methylation (Fig.3C and Fig. S6). Thus, interaction with Hbs in the absence of H-GDGTs does not allow for more robust GDGT methylation in *T. kodakarensis* suggesting that the H-GDGT methylase prefers the H-GDGT substrate over an un-bridged GDGT substrate.

### H-GDGT biosynthesis proteins are found in diverse archaea and bacteria

With the knowledge that Hbs and Hgm synthesize H-GDGTs and methylated H-GDGTs, respectively, we next sought to determine the distribution of these proteins in microbial genomes to gauge how widespread H-GDGT biosynthesis and methylation is throughout the microbial world. To do this we searched the Joint Genome Institute (JGI) and National Center for Biotechnology Information (NCBI) databases using the Tg-Hbs and Tg-Hgm as the search queries with an e-value cutoff of 1e-80 for Hbs and 1e-65 for Hgm and a minimum sequence length of 400 amino acids. Redundant sequences were removed from the dataset, the remaining sequences were aligned, and phylogenetic trees were constructed using AhbD and Moek5 sequences as outgroups for Hbs and Hgm, respectively (Fig. 4). A well-bootstrapped clade of both Hbs and Hgm proteins was identified and satisfied the following criteria: 1) separation from outgroup sequences, 2) inclusion of all archaea known to produce H-GDGTs and methylated H-GDGTs, and 3) presence of the unique COG4001 domain for Hbs homologs.

**Figure 4:**
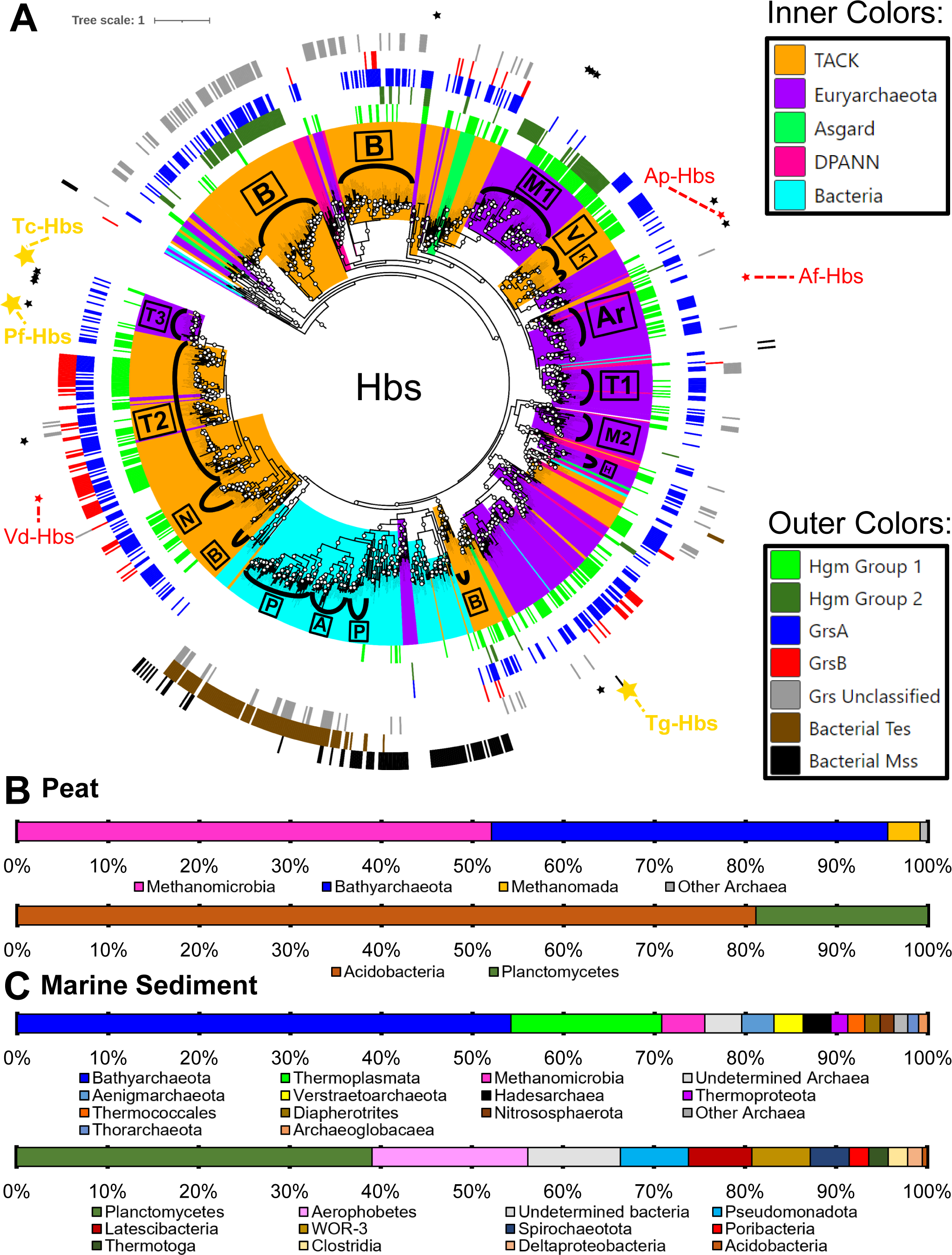
Hbs homologs are found diverse archaea and bacteria and implicate certain microbial groups as major H-GDGTs producers in the environment. A) Hbs tree showing the phylogenetic distribution of Hbs homologs and their co-occurrence with other GDGT modifying genes. Inner ring colors denotate what major phyla the Hbs sequences belong to. More specific phylogenetic clades are denoted as follows: Acidobacteria [A], Archaeoglobi [Ar], Bathyarchaeota [B], Hadesarchaea [H], Korarchaeota [K], Methanomada [M1], Methanomicrobia [M2], Nitrososphaeota [N], Planctomycetes [P], Thermoplasmata [T1], Thermoproteota [T2], Thermococcales [T3], and Verstraetearchaeota [V]. The Hbs proteins expressed in this study are denoted by gold stars, highlighting the phylogenetic difference of Tg-Hbs to Tc-Hbs and Tg-Hbs. Hbs proteins of cultured archaea that we analyzed for H-GDGT biosynthesis in this study are denoted by red stars: *A. profundus* Hbs (Ap-Hbs), *A. fulgidus* Hbs (Af-Hbs), and *Vulcanisaeta distributa* Hbs (Vd-Hbs). Other archaea which have been demonstrated to produce H-GDGTs in culture are denoted by small black stars. Colors of the outer rings correspond to the presence of other GDGT modification genes within the genomes of the Hbs encoding microbes. Note the more widespread nature of GrsA and Hgm1 homologs compared to GrsB, Grs Unclassified, and Hgm2 homologs which associate more specifically to certain clades. Also note the two distinct groups of Hbs encoding bacteria, those possessing Mss homologs and those possessing Tes homologs. B) Relative abundance of Hbs homologs clustering with specific archaeal and bacterial phyla in peat. Both archaeal and bacterial Hbs sequences are dominated by just two phyla: the Bathyarchaeota and the Methanomicrobia for archaea and the Acidobacteria and Planctomycetes for bacteria. C) Relative abundance of Hbs homologs clustering with specific archaeal and bacterial phyla in marine sediments. The Hbs sequences are found in a greater diversity of both archaea and bacteria in marine sediments compared to peat. Bathyarchaeota still dominate among the archaea, while a shift to Planctomycetes and Aerophobetes bacteria dominating occurs in marine sediments, with almost no Acidobacterial associated sequences being detected.

Utilizing this approach, we obtained 1,149 nonredundant Hbs sequences with 497 belonging to TACK archaea, 393 to Euryarchaeota, 193 to Bacteria, 39 to DPANN archaea, 23 to Asgardarchaeota, and 4 to unclassified archaea. Thus, while almost all H-GDGT producers in culture are Euryarchaeota, the potential for H-GDGT biosynthesis is more widespread than previously thought. A closer look at the archaea comprising these superphyla revealed numerous Hbs sequences belonging to Bathyarchaeota [188], Thermoprotei [145], Methanomada [88], Thermoplasmata [74], Methanomicrobia [67], Archaeoglobi [65], non-ammonia oxidizing members of the Nitrososphaerota [49], Verstraetarchaeota [40], and Thermococcales [37]. Further, many Hbs-encoding archaea also possess additional GDGT modification proteins with 448 encoding Hgm, 424 encoding GrsA, 98 encoding GrsB, and 114 encoding unclassified Grs homologs, suggesting many archaea can produce H-GDGTs with additional modifications.

Surprisingly, about 17% of the identified Hbs sequences were found in bacterial genomes (Fig. 4). More than 90% of these bacterial genomes possessing *hbs* genes also have the genes required or hypothesized to synthesize bacterial membrane spanning lipids (*mss* and/or *tes*) such as brGDGTs (Fig. S13). This co-occurrence of Hbs with Mss and/or Tes in bacterial genomes was especially striking given that all but 5 bacterial Hbs sequences are from metagenome assembled genomes (MAGs) which can be incomplete. The bacteria possessing Hbs homologs can be divided into two groups: 1) Acidobacteria [31] and Planctomycetes [71] bacteria primarily encoding Tes homologs and 2) diverse bacterial candidate phyla (Aerophobetes [11], Omnitrophota [12], WOR-3 [8], etc.) primarily encoding Mss homologs. Furthermore, the *hbs* genes throughout many diverse bacterial phyla co-occur in gene clusters containing either *tes* or *mss*, just as *hbs* and *hgm* genes often cluster together in archaea (Fig. S13). Further, the bacterial Hbs homologs have the same level of similarity to the archaeal Hbs proteins as the archaeal Hbs proteins tested in this study have to each other. For example, the *P. furiosus* Hbs is 45% identical and 63% similar to the *T. guaymasensis* Hbs (e-value = 5e-141) while the Aerophobetes bacteria Hbs is 46% identical and 64% similar to the *T. guaymasensis* Hbs (e-value = 8e-142). Altogether, these analyses suggest that the Hbs homologs found in bacterial genomes may be responsible for the formation of bridged membrane spanning lipids (e.g brH-GDGTs) in bacteria.

In similar fashion, we obtained 990 nonredundant Hgm sequences: 529 belonging to TACK archaea, 413 to Euryarchaeota, 26 to Asgardarchaeota, 15 to Bacteria, 6 to DPANN archaea, and 1 to an unclassified archaeon (Fig. S14). Interestingly, most of these sequences belong to only 3 archaeal phyla: Bathyarchaeota [307], Methanomada [254], and Thermoprotei [129]. There were smaller but notable contributions from Archaeoglobi [40], Thermoplasmata [34], Methanomicrobia [32], and Verstraetarchaeota [31], thereby expanding H-GDGT methylation beyond the three known producers in Methanomada. Furthermore, while two-thirds of *hgm*-encoding genomes also possess *hbs* genes, several do not. These include cultured archaea with complete genomes such as *Ferroglobus placidus*, *Geoglobus acetivorans*, *Methanobacterium subterraneum*, and *Methanothermobacter wolfeii*, suggesting that Hgm may function as a GDGT methylase, rather than an H-GDGT methylase, in some organisms. Additionally, many Methanomada archaea (e.g. Methanobacterium, Methanobrevibacter, Methanopyri) possess 2-3 Hgm homologs. All contain one “canonical” Hgm (termed Hgm1) that is found in clades A-C of the Hgm tree where all the known methylated H-GDGT producing archaea are found and an additional 1-2, potentially non-canonical, homologs (termed Hgm2) found in clade D (Fig. S14). Many Bathyarchaeota possess Hgm homologs exclusively in this Hgm2 clade, and the majority of the Bathyarchaeota with *hgm2* genes have multiple copies from several different areas of the Hgm2 portion of the tree: 23 with one copy, 19 with two copies, 39 with 3 copies, and 29 with 4 or more copies, some of which are found on the same metagenomic scaffold. Altogether, these analyses demonstrate that H-GDGT methylation (or GDGT methylation in some instances) is found in diverse archaea but that it is potentially more significant in Bathyarchaeota and Methanomada. These archaea frequently possess multiple Hgm homologs, some of which may have alternative activity such as different degrees or positioning of methylation or different substrate preferences.

### Investigating the biological sources of H-GDGTs in peat soils and marine sediments

Both H-GDGT and the bacterial brH-GDGT lipids are often found in peat soils and marine sediment environments, and their microbial sources in these environments have been speculated on previously (2, 30). With an H-GDGT bridge synthase protein now known, we can provide more insight into the biological sources of these lipids in the environment. To do this, we searched 2,109 peat metagenomes and 471 marine sediment genomes in the JGI database for Hbs homologs, with an e-value cut-off of 1e-80 and at least 400 amino acid sequence length. We then aligned these sequences with those previously used to make the Hbs tree and constructed phylogenetic trees using AhbD as an outgroup. In total 1,712 sequences from the peat metagenomes and 827 sequences from the marine sediment metagenomes clustered within the Hbs portion of our tree. We then assigned the metagenomic sequences to a particular taxonomic group based upon the taxonomy of the known Hbs sequences they clustered with (Fig 4B and 4C).

Based on this analysis, we found that 77% of the total Hbs peat sequences belonged to archaea, and that these archaeal sequences were dominated by just two phyla: Methanomicrobia (51%) and Bathyarchaeota (44%) (Fig. 4B). Further, >98% of the Methanomicrobia sequences clustered closely with an Hbs sequence from the methanogen *Methanoflorens stordalenmirensis* isolated from a peatbog in Sweden. These closely related Methanomicrobia sequences were found in peat environments distributed throughout the globe including those in Alberta (Canada), Barrow (Alaska), Department of Meta (Colombia), Houghton (Michigan), Ithaca (New York), Loreto (Peru), Marcell Experimental Forest (Minnesota), and Stordalen Mire (Sweden). *M. stordalenmirensis* possesses an *hgm* gene (particularly *hgm2*) immediately adjacent to *hbs* and multiple related metagenomic sequences also possess adjacent *hgm* genes, suggesting these archaea may be responsible for methylated H-GDGT production in peat environments as well. Approximately 23% of the total peat sequences belonged to bacteria and were dominated by just two phyla: 81% Acidobacteria and 19% Planctomycetes bacteria. The Acidobacterial sequences were found in all the aforementioned sites except for Barrow, Alaska while Planctomycetes Hbs homologs were slightly more restricted, being found only at the Alberta, Houghton, Marcell Experimental Forest, and Stordalen Mire sites. No cultured bacteria are known to produce brH-GDGTs, but Acidobacteria are known to produce brGDGTs and have been suggested as a source of brH-GDGTs in peat soils (30). However, the involvement of Planctomycetes bacteria in brH-GDGT production in peat environments would represent a novel source.

In marine sediments, we found that 77% of the identified Hbs homologs belonged to archaea (Fig. 4C). While these marine sediments possess a more diverse assemblage of potential H-GDGT producers than in peat metagenomes, they are still dominated by Bathyarchaeota (54%) with smaller contributions from Thermoplasmatota (15%), Methanomicrobia (4.5%), Hadesarchaea (3.4%), Aenigmarchaeota (3.3%), and Verstraetarchaeota (3.0%), amongst others (Fig. 4C). These Bathyarchaeota sequences are present in a diversity of marine sediment environments and localities including Aarhus Bay (Denmark), coastal sediments offshore of San Francisco (California), hydrothermal sediments at Pescadero Basin (Gulf of California), Little Sippewissett salt marsh (Massachusetts), the Washington margin (Washington), and White Oak Estuary (North Carolina). We also found that potential bacterial brH-GDGT producers (Fig. 4C) are also more diverse in marine sediments and are dominated by Planctomycetes (39%) and Aerophobetes (17%) bacteria with smaller contributions from Pseudomonadota (7.5%), Latescibacteria (7.0%), and WOR-3 (6.4%) phyla amongst others. Only 1 Acidobacteria- associated sequence was recovered from the 471 marine metagenomes searched, suggesting Acidobacteria are not a major source of marine sediment derived brH-GDGTs. Recently, Planctomycetes bacteria have been implicated as a potential source of highly methylated brGDGTs, termed overly branched GDGTs (obGDGTs) in Mariana Trench sediments, implying that they could be a source of “H-obGDGTs” which have not yet been detected (48) . Notably, Planctomycetes Hbs sequences are primarily found in mesophilic environments such Aarhus Bay, the Caspian Sea, coastal sediments offshore of San Francisco, Little Sippewissett salt marsh, the Washington margin, and White Oak Estuary while Aerophobetes sequences are found in some mesophilic environments (primarily Aarhus Bay and the Washington Margin) as well as higher temperature environments like the hydrothermal sediments in Pescadero Basin. Some Aerophobetes MAGs from hydrothermal vents possess GDGT ring synthase genes adjacent to *hbs*, suggesting these bacteria may produce bacterial membrane spanning lipids with both a bridge and a ring(s) which have not yet been observed (Fig. S13). Many of the other bacterial candidate phyla such as Latescibacteria, Omnitrophota, Poribacteria, Thermotogota, and WOR-3 were primarily found in hydrothermal sediments with the notable exception of the Pseudomonadota which were only found in White Oak Estuary. Altogether, these analyses suggest a major Planctomycetes contribution to the brH-GDGTs found in mesophilic marine sediments but a diverse bacterial candidate phyla contribution of high temperature marine sediment derived bacterial H-shaped lipids.

### *Archaeoglobus* archaea synthesize H-GDGTs and increase their relative abundance at elevated temperatures

While we see that the genes responsible for H-GDGT biosynthesis and methylation are widespread in genomic databases, the number of archaea known to produce these lipids in culture is small (n = 22 for H-GDGTs and n = 3 for methylated H-GDGTs) (18, 49). Therefore, we aimed to expand the diversity of cultured H-GDGT producers and to begin investigations into parameters governing the production and modification of these lipids. To do this, we analyzed the lipidomes of *Archaeoglobus fulgidus* and *Archaeoglobus profundus* in response to changes in temperature as both archaea possess an Hbs homolog and have been reported to produce “unknown” lipids consistent with H-GDGTs (50, 51). Additionally, *A. profundus* contains an Hgm homolog and two closely related GrsA homologs, one of which is found in a gene cluster with *hgm,* while *A. fulgidus* lacks both the methylase (*hgm)* and cyclization (*grs)* genes. Cultures of *A. fulgidus* and *A. profundus* were grown at 60°C, 70°C, 80°C, and 85°C and at 70°C, 80°C, and 90°C, respectively, to stationary phase. The biomass was then acid hydrolyzed, and lipids were extracted and analyzed by LC-MS in normal phase.

Robust H-GDGT production was identified in both *A. fulgidus* and *A. profundus*, and both archaea increased the relative abundance of H-GDGT lipids in response to elevated growth temperatures (Fig. 5 and Fig. S15). H-GDGTs in *A. fulgidus* increased from 0.1% of monolayer lipids at 60°C to 43% at 85°C, and H-GDGTs in *A. profundus* went from 25% at 70°C to 92% of monolayer lipids at 90°C, the latter amongst the largest proportions of H-GDGTs relative to other monolayer lipids observed in cultured archaea (18, 49) . We additionally detected H-GDGT methylation in *A. profundus*, expanding known methylated H-GDGT producers beyond thermophilic methanogens in Methanomada. We found similar levels of methylation at 70°C and 80°C, with 9.8% and 7.0% of H-GDGTs being mono-methylated, respectively, and a slightly higher degree of methylation (16%) at 90°C. Di-methylated H-GDGTs were not detected and only trace GDGT methylation (at most 0.4% of GDGTs) was observed.

**Figure 5.**
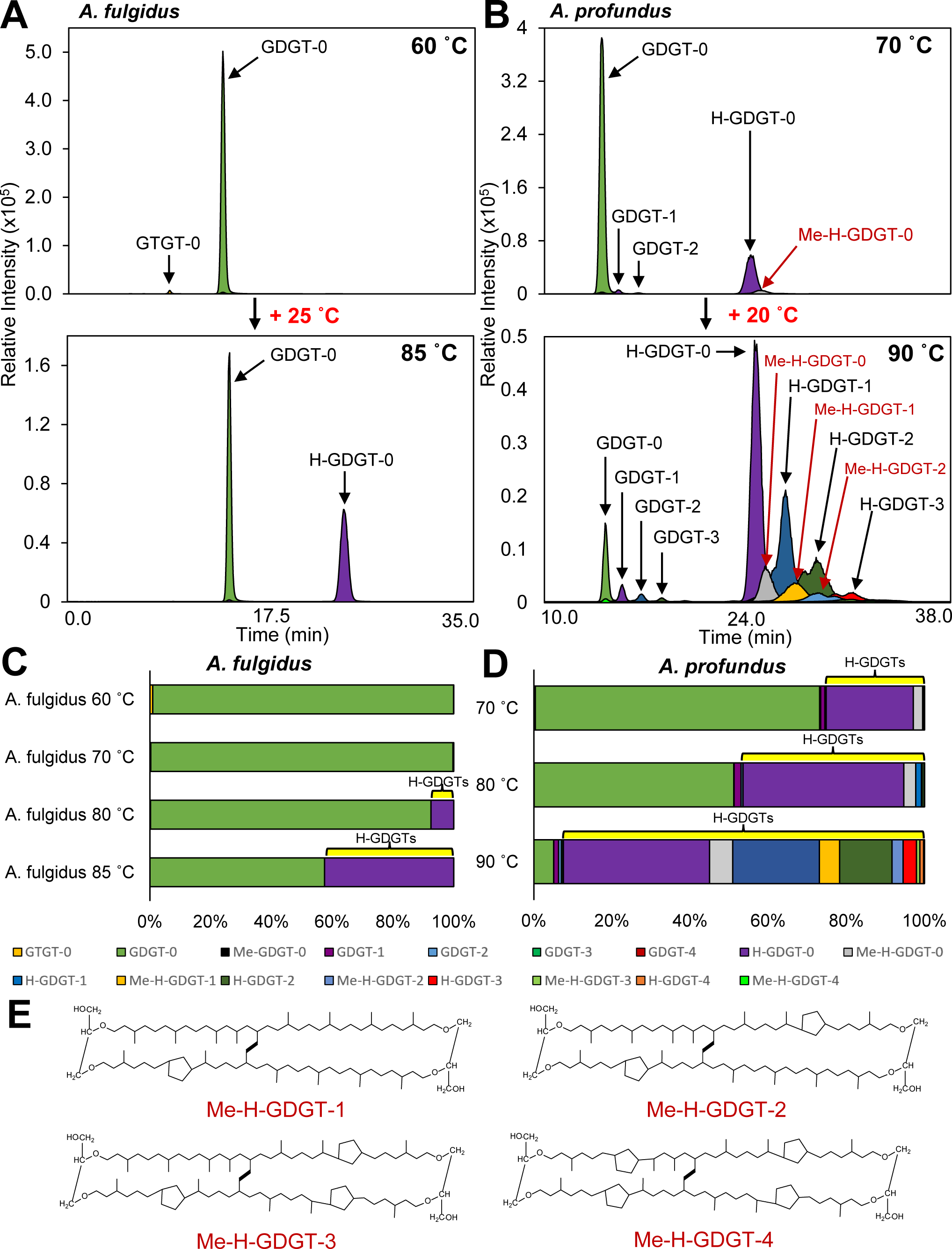
*A. fulgidus* and *A. profundus* produce H-GDGTs and increase their relative abundance in response to elevated growth temperatures. A) Merged EICs of *A. fulgidus* lipids showing the differences in core lipids produced during growth at 60°C and 85°C. Abundant GDGT-0 and small amounts of GTGT-0 are seen at 60°C while abundant GDGT-0 and abundant H-GDGT-0 can be seen at 85°C. B) Merged EICs of *A. profundus* lipids showing the differences in core lipids produced during growth at 70°C and 90°C. Abundant GDGT-0 and small amounts of ring containing GDGTs are seen at 70°C along with moderate amounts of H-GDGT-0 and a small amount of methylated H-GDGT-0. A large diversity of core lipids structures is produced at 90°C including GDGTs and their cyclized derivatives and H-GDGTs and their cyclized and methylated derivatives. C) Relative abundance of monolayer lipids in *A. fulgidus* across temperature, showing that H-GDGT production is stimulated between 70°C and 80°C and gradually increases with temperature. D) Relative abundance of monolayer lipids in *A. profundus* across temperature, showing that H-GDGTs gradually increase in their relative abundance with increased growth temperature, eventually becoming dominant over GDGTs and accounting for >90% of total monolayer lipids. E) Monolayer lipids from *A. profundus* which have not been observed in culture prior to this study. These triply modified GDGT- type lipids possess a bridge, a methylation, and rings and are most abundant at the highest growth temperatures.

We also detected both GDGT and H-GDGT cyclization in *A. profundus*, observing up to 4 rings in both core structures depending on the temperature, and seeing an increase in the average number of rings per molecule (the ring index or RI) in response to temperature (Fig. 5). The ring index shifted from an RI_GDGT_ and an RI_H-GDGT_ of 0.02 and 0.02 at 70°C, respectively, to an RI_GDGT_ and an RI_H-GDGT_ of 0.7 and 0.9 at 90°C, respectively. For the first time in culture, we detect the presence of triply modified GDGT- based lipids – that is Me-H-GDGT-1, Me-H-GDGT-2, Me-H-GDGT-3, and Me-H-GDGT-4 which contain 1) a bridge/cross-link, 2) a methylation, and 3) cyclopentane rings (Fig. 5). Such lipids are present in only trace amounts at 70°C and 80°C (at most a combined 0.2% of monolayer lipids) but make up 9.1% combined at 90°C with Me-H-GDGT-1 and Me-H- GDGT-2 being the major triply modified lipids. Further, the degree of methylation of the different cyclized H-GDGTs and acyclic H-GDGT were all similar, ranging from 14-20% at 90°C, suggesting that either the presence of rings does not affect the activity of Hgm or that the presence of methylations does not affect Grs activity in *A. profundus*. Despite these additional modifications and different lipid biosynthetic gene inventories, we see that H-GDGTs appear to play an important role in growth at elevated temperatures in both *A. fulgidus* and *A. profundus.* This could be a common feature of the genus Archaeoglobus more broadly as the other members of this group with available genomes (*Archaeoglobus neptunis*, *Archaeoglobus sulfaticallidus*, and *Archaeoglobus veneficus*) all possess Hbs homologs as do many MAGs belonging to this genus.

### Investigating the biological sources of ARN acid precursors

H-GDGTs can possess additional modifications including methylations and rings such as those seen in this study and previous works in culture and the environment (18, 19, 25, 26). However, highly cyclized H-GDGTs, particularly H-GDGT-5 and H-GDGT-6 possessing 5 and 6 cyclopentane rings, respectively, have so far been solely identified in hot spring environments and have not been observed in culture (29). Cyclized H-GDGTs are of additional interest as they are the likely precursors of a group of undesirable compounds in crude oil - the ARN acids (Fig. 6). ARN acids are the oxidative breakdown products of H-GDGTs and most commonly possess 6 cyclopentane rings when found in crude oil (35). To produce H-GDGT-5 and -6, microbes require three GDGT modification genes: 1) Hbs to cross-link the tails, 2) GrsA to add 4 cyclopentane rings at the C-7 positions along the molecule, and 3) GrsB to add 1-2 cyclopentane rings at C-3 positions in the molecule (Fig. 6A). As such, we searched the genomes of all Hbs-encoding archaea for the presence of both GrsA and GrsB homologs. In total, we identified 76 archaeal genomes, 11 from cultured archaea and 65 MAGs, possessing the *hbs*, *grsA*, and *grsB* genes (Fig. 6B and Supp. Data Set 1). Most of these genomes belong to archaea within Thermoprotei (n = 54) with smaller contributions from various Euryarchaeota (n = 8), Bathyarchaeota (n = 7), Nitrososphaerota (n = 6), and Odinarchaeota (n = 1), revealing diverse phyla are capable of highly cyclized H-GDGT production.

**Figure 6.**
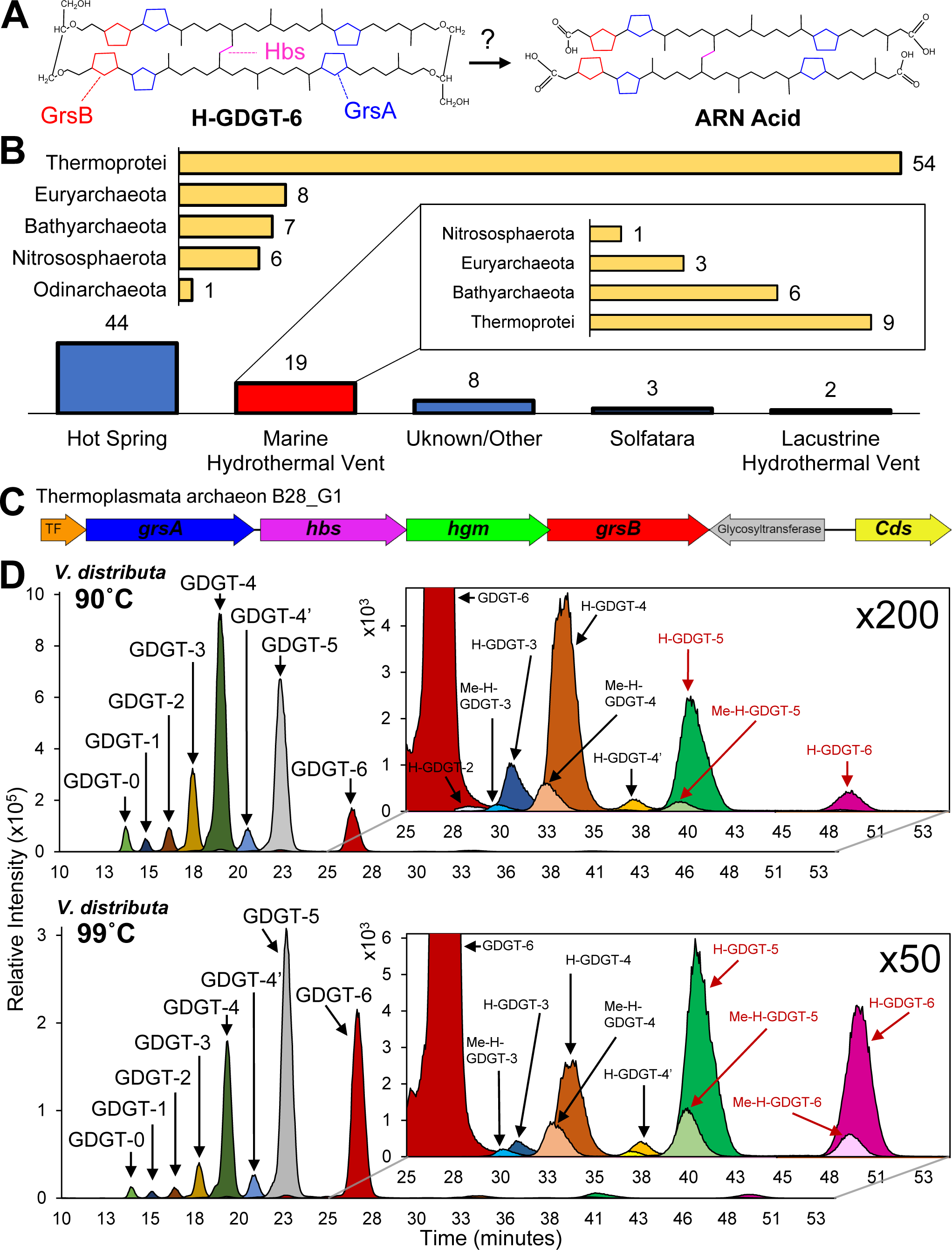
Potential highly cyclized H-GDGT producers are diverse yet distinctive. A) Structures of the highly cyclized H-GDGT-6 and its presumed ARN acid derivative in crude oil. GrsA, GrsB, and Hbs are all required to synthesize the modifications found in such lipids. B) Phylogenetic and environmental distributions of highly cyclized H-GDGT candidate producers identified in this study. Genomes belonging to Thermoprotei archaea dominate as do genomes from hot springs. The phylogenetic distribution of potential highly cyclized H-GDGT producers in marine hydrothermal vent ecosystems is not as strongly dominated by the Thermoprotei, suggesting distinct sources of these lipids in these two different environments. C) Gene cluster containing all of the GDGT modification genes necessary to synthesize highly cyclized H-GDGTs in a Thermoplasmata archaeon MAG from Guaymas Basin. TF = transcription factor, Cds = calditol synthase. D) Merged EICs showing the large diversity of core lipids present in *V. distributa* during growth at 90°C and 99°C. GDGTs dominate the monolayer lipids present in *V. distributa*, but H-GDGT-5 and H-GDGT-6, along with their methylated derivatives, were seen for the first time here in a cultured archaeon.

Interestingly, most of these genomes (84%) also encoded Hgm homologs indicating that these archaea might also be methylating their highly cyclized H-GDGTs. As expected, the majority of these MAGs and cultures originated from hot springs (58%), with marine hydrothermal vents being the second most common environment (26%). Hot springs are a very different environment from oilfields (sources of ARN acids), and thus the H-GDGT- 5 and -6 producers in these locations may very well differ. While there are few available archaeal isolates or MAGs from oilfield/reservoir sites (n = 10 in JGI), we identified one archaeon, *Thermoplasma kamchatkensis* strain Kam2015, that was isolated from an oilfield in Russia that possesses *hbs*, *grsA*, and *grsB* genes. Thus, archaea with the genetic potential to produce highly cyclized H-GDGTs do inhabit environments where crude oil is present and suggests these producers may be distinct from the Thermoprotei, which appear to be the dominant producers of highly cyclized H-GDGT in hot springs.

Marine hydrothermal vent systems are often also rich in oil and ARN acids have been detected in hydrothermal vent sediments from Guaymas Basin while being notably absent from hot springs, even though both localities possess cyclized H-GDGTs (35). As such, we performed a more in-depth analysis of the 19 MAGs from the marine hydrothermal vent (mostly sediment) ecosystems: 9 from Thermoprotei, 6 from Bathyarchaeota, 3 from Euryarchaeota, and 1 from Nitrososphaerota. Interestingly, most of these MAGs, especially those outside Thermoprotei, were identified in the oil rich sediments of Guaymas and Pescadero Basin. Of particular interest are two Euryarchaeota MAGs within Alkanophaga and Thermoplasmata. The first MAG belongs to *Candidatus* Alkanophaga volatiphilum, an “oil eating” (C_5_-C_7_ hydrocarbons) microbe identified in an anoxic, thermophilic (70 °C) enrichment culture from Guaymas Basin sediments (52). It is plausible that organisms like *Candidatus* Alkanophaga volatiphilum would associate closely with oil in these settings and their highly cyclized H-GDGTs would be deposited and diagenetically degraded into ARN acids. Alternatively, these archaea may enzymatically degrade their H-GDGTs, biotically producing the ARN acids to use as biosurfactants to aid in the utilization and consumption of the oil as suggested by Lutnaes et al. (16). Other hydrocarbon degrading archaea, such as *Alkanophaga liquidiphilum*, *Methanolliviera hydrocarbonicum*, and *Syntrophoarchaeum butanivorans*, possess both Hbs and GrsA homologs and could therefore produce moderately cyclized H-GDGTs with up to 4 rings. While ARN acids with 5, and especially 6, rings are the most abundant in most environmental sites analyzed, those with 4 rings are often the next most abundant and have been shown to dominate in some locations (17, 53–55). Thus, these hydrocarbon degrading archaea that generate moderately cyclized H-GDGTs may also play a role, albeit more minor, in ARN acid production.

A second MAG, Thermoplasmata archaeon isolate B28_G1, also from Guaymas Basin, stands out due to the clustering of the *grsA*, *hbs*, *hgm*, and *grsB* GDGT modification genes in its genome in an operon-like structure (Fig. 6C). The placement of these genes suggests that they may be co-regulated or co-expressed, and the products of such expression would theoretically result in the synthesis of highly cyclized, methylated H-GDGTs some of which have been detected in Guaymas Basin (35). We additionally found multiple B28_G1 related metagenomic scaffolds also possessing co-organized *grsA*, *hbs*, *hgm*, and *grsB* genes in JGI metagenomes from the hydrocarbon rich sediments of Guaymas and Pescadero Basin (Fig. S16A), further corroborating the role of these archaea in H-GDGT-5 and -6 production at such localities. Taken together, these data suggest that the Thermoplasmata group is one likely source of ARN acids in crude oil as they have the biosynthetic potential to produce highly cyclized H-GDGTs, organize their GDGT biosynthesis genes for the production of such lipids, and occur in multiple localities (Uzon oilfield, Guaymas and Pescadero basin) that are rich in hydrocarbons or oils.

### Highly cyclized H-GDGT production and methylation in a Thermoproteales archaeon

Having identified a handful of cultured archaea that possess the genetic potential to produce highly cyclized H-GDGTs, we next wanted to confirm production of these lipids in culture. Besides *T. kamchatkensis*, all the cultured archaea encoding Hbs, GrsA, and GrsB are anaerobic thermoacidophiles isolated from hot springs and consist of 9 Thermoprotei archaea, particularly in Acidilobales and Thermoproteales, and 1 Nitrososphaerota archaeon, *Conexivisphaeara caldida*. Of these 11 candidate H-GDGT- 5 and 6 producing isolates, we selected the Thermoproteales archaeon *Vulcanisaeta distributa* for further analysis. Previous studies (56) have investigated the lipid composition of *V. distributa* and reported only the presence of GDGT-0 and various cyclized GDGTs. Through our lipid analysis, we also found that GDGT-0 to GDGT-6 were the dominant (∼99%) monolayer lipids in *V. distributa* grown at optimal conditions (90°C and pH = 4.25) with GDGT-4 being the most abundant (Fig. 6D and Fig. S16B). However, we were also able to detect small amounts of H-GDGT production (0.9% of monolayer lipids), detecting H-GDGT-2 to H-GDGT-6, of which H-GDGT-4 was the most dominant. Additionally, *V. distributa* possesses an Hgm homolog and we found that ∼7-14% of each H-GDGT was methylated under these conditions with H-GDGTs possessing more rings (5–6) being on the lower end of this proportion. We next sought to determine if we could stimulate H-GDGT production in *V. distributa* as in *A. fulgidus* and *A. profundus* by increasing the growth temperature. While increasing the growth temperature to 99°C did not result in abundant H-GDGT production, the relative abundance of H-GDGTs did increase to ∼4.4% with H-GDGT-5 and H-GDGT-6 being the dominant H-GDGTs under these conditions (Fig. 6D and Fig. S16B). Additionally, we noted an approximate doubling in the degree of methylation of each H-GDGT at 99°C compared to 90°C, ranging from 12-33% methylated, with the H-GDGTs containing 5-6 rings again being on the lower end of this proportion. The ring distribution of the GDGTs as in the H-GDGTs also shifted, with GDGT-5 and GDGT-6 being the dominant monolayer lipids under these conditions.

## Conclusion

The biosynthesis and physiology of the unique GDGT membrane lipids harbored by archaea have been of great interest since their discovery and characterization nearly half a century ago (57, 58). In this study, we identify two proteins responsible for the formation and methylation of one of the more distinctive GDGT lipids - the cross-linked H-GDGTs. The H-GDGT bridge synthase (Hbs), required for the cross-linking of the two alkyl chains of the core GDGT, and the H-GDGT methylase (Hgm) are both radical SAM proteins emphasizing again the broad importance of radical SAM enzymology in the synthesis and maintenance of membrane lipids in archaea. By linking bioinformatic analyses with heterologous expression, lipid analyses, and physiological characterization, we revealed potentially novel H-GDGT producers, the dominant sources in a variety of ecosystems, and a physiological link to increased temperature stress in certain archaeal species. More broadly, this study shows that the discovery and characterization of proteins required to synthesize archaeal lipids continues to be a powerful approach that allows us to explore and better understand the physiological, biochemical, and geochemical implications of these complex lipids.

## Materials and Methods

### Microbial Strains, Media, and Growth Conditions

The strains used and constructed in this study are listed in Table S1. All *Escherichia coli* strains were grown at 37°C on Lysogeny Broth (LB) supplemented with the following antibiotics as needed: 100 µg/mL Carbenicillin and 30 µg/mL Kanamycin. All *Thermococcus kodakarensis* strains were grown in anaerobically prepared artificial sea water (ASW) medium supplemented with 5 g/L tryptone, 5 g/L yeast extract, 2 g/L elemental sulfur, Wolfe’s trace elements, and KOD vitamins as described previously (59). The parental *T. kodakarensis* strain AL010 (containing no plasmid) is an agmatine auxotroph and was additionally supplemented with 1 mM agmatine sulfate. *T. kodakarensis* cells grown for lipid analyses were inoculated in a Coy anaerobic chamber at an OD_600_ = 0.01 and were cultured in 125 mL serum vials containing 100 mL of media at 85°C to mid-stationary phase (t = 14 hours). Cells were immediately harvested by centrifugation at 4500 x g at 4°C for 30 minutes and stored at –80°C until lipid extraction.

*Archaeoglobus fulgidus* Z (DSM 4139) and *Archaeoglobus profundus* AV18 (DSM 5631) were acquired from Deutsche Sammlung von Mikroorganismen und Zellkulturen (DSMZ). *A. fulgidus* Z was cultivated in media based on Zellner et al. (60) with the following components per l: KCl, 0.335g; MgCl_2_·6H_2_O, 2.75g; MgSO_4_, 1.684g; Na_2_SO_4_, 4.26g; CaCl_2_·2H_2_O, 0.14g; K_2_HPO_4_, 0.14g; NaCl, 18g; Yeast Extract, 2g; Peptone, 2g; Na-Acetate, 0.6g; Na-Pyruvate, 5g; FeSO_4_·7H_2_O (0.1% w/v), 1.36ml; NiSO_4_·6H_2_O (0.1% w/v), 1.6ml; Na_2_WO_4_·2H_2_O (0.1% w/v), 38µl; Na-resazurin (0.1% w/v), 0.5ml; Trace element solution, 10ml; Vitamin solution, 10ml; NaHCO_3_, 5g; NH_4_Cl, 0.25g; L- cysteine·HCl, 0.5g; Na_2_S·9H_2_O, 0.5g. Cells were grown for 48 hours and harvested in stationary phase by centrifugation at 5000 x g for 15 minutes at 4°C and stored at -80°C until lipid extraction. *A. profundus* AV18 was cultivated in media based on DSMZ medium 519 with the following components per l: KCl, 0.34g; MgCl_2_·6H_2_O, 4g; MgSO_4_, 1.684g; Na_2_SO_4_, 2.7g; CaCl_2_·2H_2_O, 0.14g; K_2_HPO_4_, 0.14g; NaCl, 18g; Yeast Extract, 1g; Na-Acetate, 1g; FeSO_4_·7H_2_O (0.1% w/v), 1.36ml; NiSO_4_·6H_2_O (0.1% w/v), 1.6ml; Na_2_WO_4_·2H_2_O (0.1% w/v), 38µl; Na-resazurin (0.1% w/v), 0.5ml; Trace element solution, 10ml; NaHCO_3_, 1g; NH_4_Cl, 0.25g; L-cysteine·HCl, 0.5g; Na_2_S·9H_2_O, 0.5g. Vitamin solution contains the following components per l: p-Aminobenzoic acid, 10 mg; Nicotinic acid, 10 mg; Ca- panthotenate, 10 mg; Pyridoxine·HCl, 10 mg; Riboflavin, 10 mg; Thiamine·HCl, 10 mg; Biotin, 5 mg; Folic Acid, 5 mg; α-Lipoic Acid, 5 mg; Vitamin B12, 5 mg. Trace element solution contains the following components per l: Nitrilotriacetic acid (trisodium salt), 1.5g; Fe(NH_4_)_2_(SO_4_)_2_, 0.8g; Na_2_SeO_3_, 0.2g; CoCl_2_·6H_2_O 0.1g MnSO_4_·H_2_O, 0.1g; Na_2_MoO_4_·2H_2_O, 0.1g; Na_2_WO_4_·2H_2_O, 0.1g; ZnSO_4_·7H_2_O, 0.1g; NiCl_2_·6H_2_O, 0.1 g; H_3_BO_3_, 0.01g; CuSO_4_·5H_2_O, 0.01g. All components excluding NaHCO_3_, NH_4_Cl, L- cysteine·HCl, Na_2_S·9H_2_O were added to 1L of water and sparged with N_2_:CO_2_ (80:20) gas mix for thirty minutes to remove oxygen. Media was then brought into a Coy chamber where solid L-cysteine·HCl was added. Once resazurin cleared NaHCO_3_, NH_4_Cl and Na_2_S·9H_2_O were added. 500ml media was dispensed in 1L bottles in the anaerobic chamber, stoppered, and headspace was made 10psi overpressure N_2_:CO_2_ (80:20) gas mix before autoclaving. For *A. profundus* AV18 cultivation headspace was 30psi overpressure H_2_:CO_2_ (80:20) gas mix. Cells grown for 72 hours and harvested in stationary phase by centrifugation at 5000 x g for 15 minutes at 4°C and stored at -80°C until lipid extraction.

*Vulcanisaeta distributa* DSM 14429 was acquired from DSMZ and was cultured in anaerobically prepared Brock medium containing the following components per liter: 1.30 g (NH_4_)_2_SO_4_, 0.28 g KH_2_PO_4_, 0.25 g MgSO_4_ x 7H_2_O, 0.07 g CaCl_2_ x 2H_2_O, 0.012g/L FeCl_3_, and 5 mL of 2000X Allen’s trace element solution. The media was supplemented with 0.5 g/L yeast extract, 0.1% dextrin from maize starch, and 1.0 g/L sodium thiosulfate as an electron acceptor and was reduced with 0.5 g/L Na_2_Sx9H_2_O. The yeast extract, dextrin, FeCl_3_, trace minerals, and Na_2_Sx9H_2_O were added after autoclaving, and the pH was adjusted to 4.25 after adding Na_2_Sx9H_2_O by the addition of sulfuric acid. *V. distributa* was cultivated anaerobically at 90 °C or 99 °C in 125 mL serum vials containing 80 mL media for 4 days to stationary phase. Cells were harvested by centrifugation at 4500 x g at 4°C for 30 minutes and stored at –80°C until lipid extraction.

### Molecular Cloning and Strain Construction

All primers used in this study are listed in Table S2. Hbs genes from *Thermococcus guaymasensis* DSM 11113 (Locus tag: X802_RS00105), *Thermococcus celer* Vu13 (Locus tag: A3L02_02305), and *Pyrococcus furiosus* (Locus tag: PF0647) were synthesized with the strong promoter pcsg (44) immediately upstream of the start codon and cloned into the self-replicative plasmid pTS543 at the NotI site. PF0647 (Pf- *hbs*) was codon optimized for expression in *T. kodakarensis* while X802_RS00105 and A3L02_02305 were not, having codon adaptation index values similar to the genes of *T. kodakarensis*. For the *T. guaymasensis* Hbs + Hgm co-expression plasmid, X802_RS00105 and X802_RS00110 were synthesized together with one pcsg promoter immediately upstream of the start codon of Tg-*hbs*. The Tg-*hbs* and Tg-*hgm* sequences overlapped exactly as they do in *T. guaymasensis* (X802_RS00105 naturally possesses a ribosome binding site for X802_RS00110). To construct the Hgm expression plasmid, the pTS543-pcsg-*hbs*-*hgm* co-expression plasmid was amplified by polymerase chain reaction (PCR) with primers flanking both ends of the *hbs* gene to amplify the other regions of the plasmid, subsequently digested with DpnI, and transformed into *E. coli* DH10B. Site-directed mutagenesis to convert the CXXXCXXC iron-sulfur binding motif to AXXXAXXA was achieved by PCR with a single primer containing the desired nucleotide changes followed by DpnI digestion and transformation into *E. coli* DH10B.

*T. kodakarensis* strains were constructed based on agmatine auxotrophy as described in Liman et al. (59). Briefly, a 100 mL culture of the parental *T. kodakarensis* strain AL010 was grown to early stationary phase (10-12 hours) to reach competence and was then anaerobically transferred to 50 mL conical tubes and spun at 4500 x g at 4°C for 30 minutes. Cells were washed and resuspended in 1.5 mL of 0.8X ASW. 200 µL aliquots of cell suspension were transferred to 1.5 mL conical tubes, placed on ice, and 2-3 µg of the desired plasmid DNA was added. Cells were incubated for 30 minutes on ice and then heat shocked for 45-60 seconds at 85°C, and then returned to ice for 30 additional minutes. Following this, 100 µL of transformed cells were plated in the absence of agmatine on ASW + Gelzan (10 g/L) containing 5 g/L tryptone, 5 g/L yeast extract, 2 mL/L of a polysulfide solution made from combining 10 g Na_2_Sx9H_2_O with 3 g of elemental sulfur in 15 mL of DI water, Wolfe’s trace minerals, and KOD vitamins. Cells were sealed inside a metal container kindly provided by the Santangelo lab at Colorado State University and incubated at 85°C for 3 days after which single colonies were picked. The presence of pTS543 and the correct insert sequence were confirmed by PCR.

### Lipid Extraction and Analyses

Frozen cell pellets were thawed and resuspended with 2mL methanol (MeOH), transferred to glass vials, and solvent was evaporated under a N_2_ stream. Samples were then acid hydrolyzed in 2 mL 1 M hydrochloric acid (HCl) n MeOH within sealed vials for 3 hours at 95°C. The reaction was neutralized by the addition of 1 mL 2M potassium chloride (KCl) in MeOH and diluted with 5 mL DI water. Lipids were then extracted three times with 5 mL of dichloromethane (DCM) which was pooled and evaporated off under a N_2_ stream. *T. kodakarensis*, *A. profundus*, and *A. fulgidus* lipid extracts were resuspended in 1 mL of 99:1 hexane:isopropanol for normal phase (NP) chromatography or 9:1 MeOH:DCM for reverse phase (RP) chromatography. *V. distributa* biomass and lipid extracts were less than the other strains and were resuspended in 100 uL of 99:1 hexane:isopropanol for NP chromatography. The resuspended lipids were then filtered through 0.45 µm polytetrafluoroethylene (PTFE) filters into fired and solvent washed glass LC-MS vials.

Lipids were analyzed via liquid chromatography-mass spectrometry (LC-MS) on an Agilent 1260 Infinity II series high performance liquid chromatography (HPLC) instrument coupled to an Agilent G6125B single quadrupole mass spectrometer using atmospheric pressure chemical ionization (APCI) in positive mode with a drying gas temperature of 250 °C, drying gas flow rate of 6.0 L/min, nebulizer pressure of 50 psi, capillary voltage of 3000V, fragmentor voltage of 175V, and a corona current of 5.0 µA. For quantification, samples were analyzed in single ion monitoring mode (SIM) for a combination of one or more of the following mass to charge (*m/z*) values when appropriate: 1342 (3-Me-H- GDGT-0), 1330 (2-Me-GDGT-0), 1328 (2-Me-H-GDGT-0), 1316 (Me-GDGT-0), 1314 (Me-H-GDGT-0), 1312 (Me-H-GDGT-1), 1310 (Me-H-GDGT-2), 1308 (Me-H-GDGT-3), 1306 (Me-H-GDGT-4 and 4’), 1304 (GTGT-0 and Me-H-GDGT-5), 1302 (GDGT-0 and Me-H- GDGT-6), 1300 (GDGT-1 and H-GDGT-0), 1298 (GDGT-2 and H-GDGT-1), 1296 (GDGT- 3 and H-GDGT-2), 1294 (GDGT-4, GDGT-4’, and H-GDGT-3), 1292 (GDGT-5, H-GDGT-4, and H-GDGT-4’), 1290 (GDGT-6 and H-GDGT-5), and 1288 (H-GDGT-6), corresponding to the [M+H]^+^ ions of the compounds listed in parenthesis. Total ion chromatograms (TICs) were also collected for all strains in scanning mode with m/z range of 600-1400.

For normal phase chromatography, lipids were separated on two Prevail Cyano columns (150 x 2.1 mm, 3 µm) connected in series with a mobile phase A of 99:1 hexane:ispropanol and a mobile phase B of isopropanol, following a method modified from (61). A flow rate of 0.3 mL/min was used with the following gradient: start with 100% A and hold for 35 minutes, linearly ramp to 93A:7B over 7 minutes, then to 90A:10B over 3 minutes, 30A:70B over 5 minutes, back to 90A:10B over 10 minutes, and back to 100% A over 10 minutes, for a total runtime of 85 minutes and with an equilibration time of 15 minutes between runs. For reverse phase chromatography, a Kinetex 1.7 µm XB-C18 100 Å LC column (150 x 2.1 mm) was used with a mobile phase A of MeOH and a mobile phase B of isopropanol following a method based off Rattray and Smittenberg (62). A flow rate of 0.2 mL/min was used with the following gradient: start with 60A:40B and hold for 1 minute, linearly ramped to 50A:50B over 19 minutes and hold this composition for 15 minutes, then return to 60A:40B over 10 minutes, with a total runtime of 45 minutes and with an equilibration time of 10 minutes between runs. An injection volume of 1 uL was used for all strains in both NP and RP chromatography, except for *A. profundus* and *V. distributa*, where 5 and 10 uL injections were used, respectively.

To obtain MS/MS spectra for 2Me-H-GDGT-0, Me-H-GDGT-0, Me-GDGT-0, and H-GDGT-0, select samples were run on a Waters Acquity UPLC and Thermo Exploris 240 BioPharma Orbitrap mass spectrometer at the Stanford University Mass Spectrometry (SUMS) facility with an APCI interface. Full scan MS spectra were collected in positive ion mode with a mass range of 200-1700 Da and Resolution 120000. Data dependent product ion spectra were collected for the top 4 most intense ions in the full scan with an intensity cutoff of 5e^3^ and absolute HCD collision energy of 40. Separation of lipids was achieved in reverse phase using a method modified from Rattray and Smittenberg (62) with a mobile phase A of MeOH and mobile phase B of MeOH. A flow rate of 0.43 mL/min was used with the following gradient: start with 60A:40B and hold for 1 minute, linearly ramp to 50A:50B over 9 minutes, hold this composition for 4 minutes, return to 60A:40B over 1 minute, and hold this composition for 5 minutes, for a total runtime of 20 minutes. Compounds were identified by their characteristic tandem mass spectral fragmentation patterns and comparison to published spectra (25, 26, 63).

### Bioinformatic and Phylogenetic Analyses

The *T. guaymasensis* Hbs (WP_062369926.1) and Hgm (WP_062369928.1), *Sulfolobus acidocaldarius* GrsA (saci_1585), *T. kodakarensis* Tes (TK2145), *Thermoanaerobacter ethanolicus* Mss (WP_129545148.1), *M. barkerii* AhbD (MA0573), and *Streptomyces viridosporous* Moek5 (WP_026084738.1) proteins were used as queries for the BLASTP (64) searches of each respective protein in the NCBI (https://www.ncbi.nlm.nih.gov/) and JGI Integrated Microbial Genomes & Microbiomes (https://img.jgi.doe.gov/) databases. For the phylogenetic analyses of Hbs and Hgm, redundant sequences were removed using CD-hit (clustered at 100% sequence identity) (65) and the remaining sequences were then combined with nonredundant AhbD or Moek5 outgroup sequences, respectively. The combined sequences were aligned using MAFFT on XSEDE (7.505) (66) and the resulting alignments were used to construct phylogenetic trees with FastTreeMP on XSEDE (2.1.10) (67). Trees were visualized using the Interactive Tree of Life (iTOL) (68).

Grs homologs were identified by searching the NCBI and JGI databases with an e-value cutoff of 1e-20 and length cutoff of >/= 400 amino acids. The obtained sequences were then used to construct phylogenetic trees as above. Clades of Grs homologs were classified as being either a GrsA, GrsB, or an unclassified Grs based on 1) the characterized Grs homologs (if present) found within each clade and 2) the published lipid profiles of archaea found within each clade. The utility of this approach to correctly classify Grs homologs is corroborated by our finding that *Vulcanisaeta distributa* can synthesize GDGT-5 and 6 – *V. distributa* possesses two Grs homologs, one that is a clear GrsA (74% similarity and 54% identity with an e-value = 0) and a second that is not clearly a GrsA or GrsB based on similarity alone, having 55% similarity and 34% identity (e-value = 5e-74) with GrsA and 50% similarity and 30% identity (e-value = 5e-65) with GrsB. However, with our approach we correctly classified this Grs homolog as a GrsB as we found that the clade which this second *V. distributa* Grs homolog belongs to has archaea that have been shown to produce GDGTs with more than 4 cyclopentane rings (i.e. capable of cyclization at C-3) while lacking close homologs to the *S. acidocaldarius* GrsB.

Tes homologs were identified by searching the NCBI and JGI databases with an e-value cutoff of 1e-40 and length cutoff of >/= 400 amino acids. The obtained sequences were then used to construct phylogenetic trees as above. To be as strict as possible, only bacterial sequences branching within the archaeal portion of the tree were classified as true Tes homologs.

Mss homologs were identified by searching the NCBI and JGI databases with an e-value cutoff of 1e-50 and length cutoff of >/= 400 amino acids. All sequences meeting these criteria were classified as Mss homologs.

## Acknowledgements

We would like to thank the Santangelo lab at Colorado State University for the generous gift of the *Thermococcus kodakarensis* AL010 strain, cuboid, and their expertise, guidance, and helpful discussions. We would also like to thank the Dekas and Francis labs at Stanford University for the use of their anaerobic chambers and members of the Welander lab for helpful discussion. This work was supported with funding from the Moore–Simons Project on the Origin of the Eukaryotic Cell Award 735931 to PVW.

## Supplemental Information

**Table S1.**
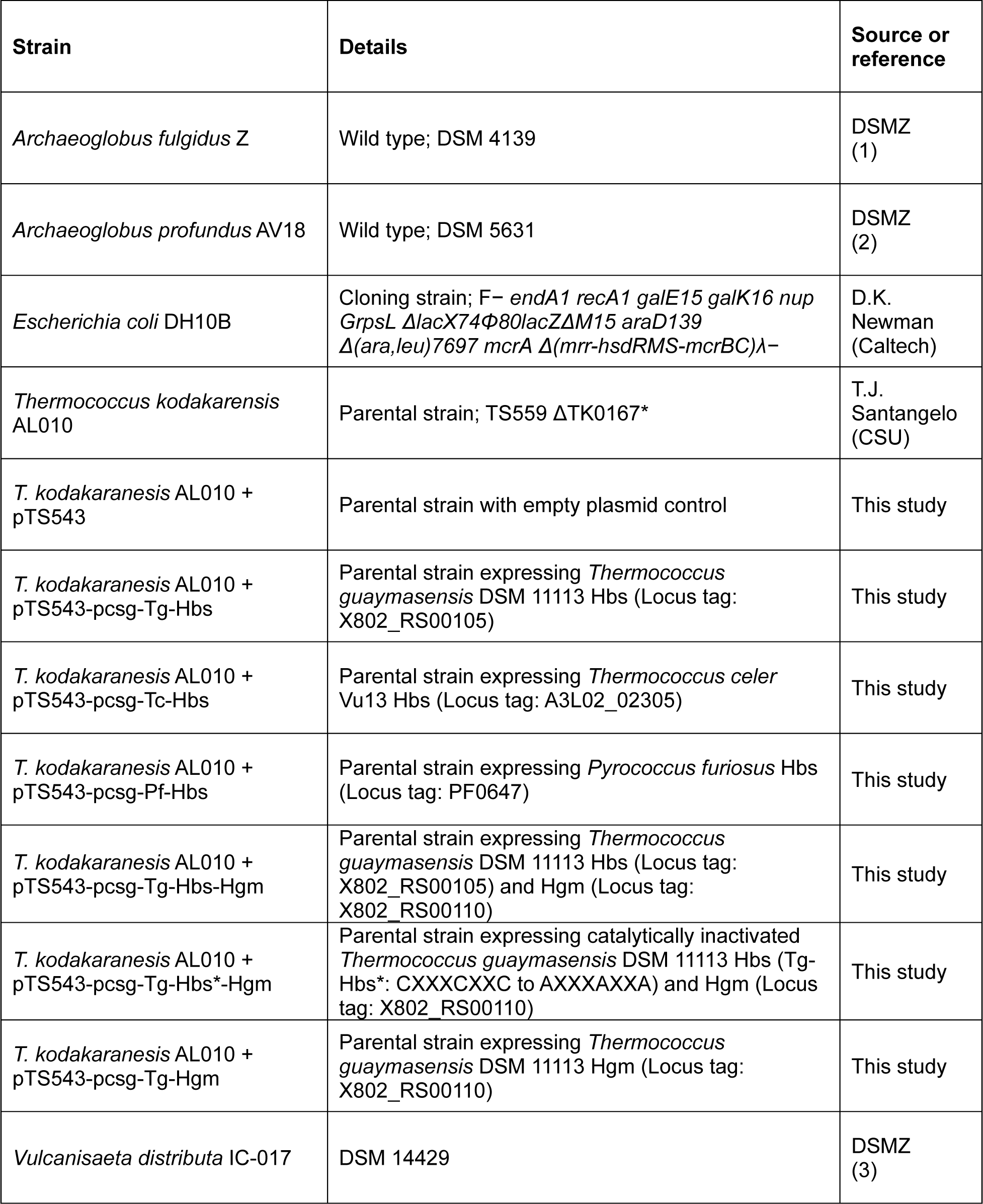
Strains used in this study.

**Table S2.**
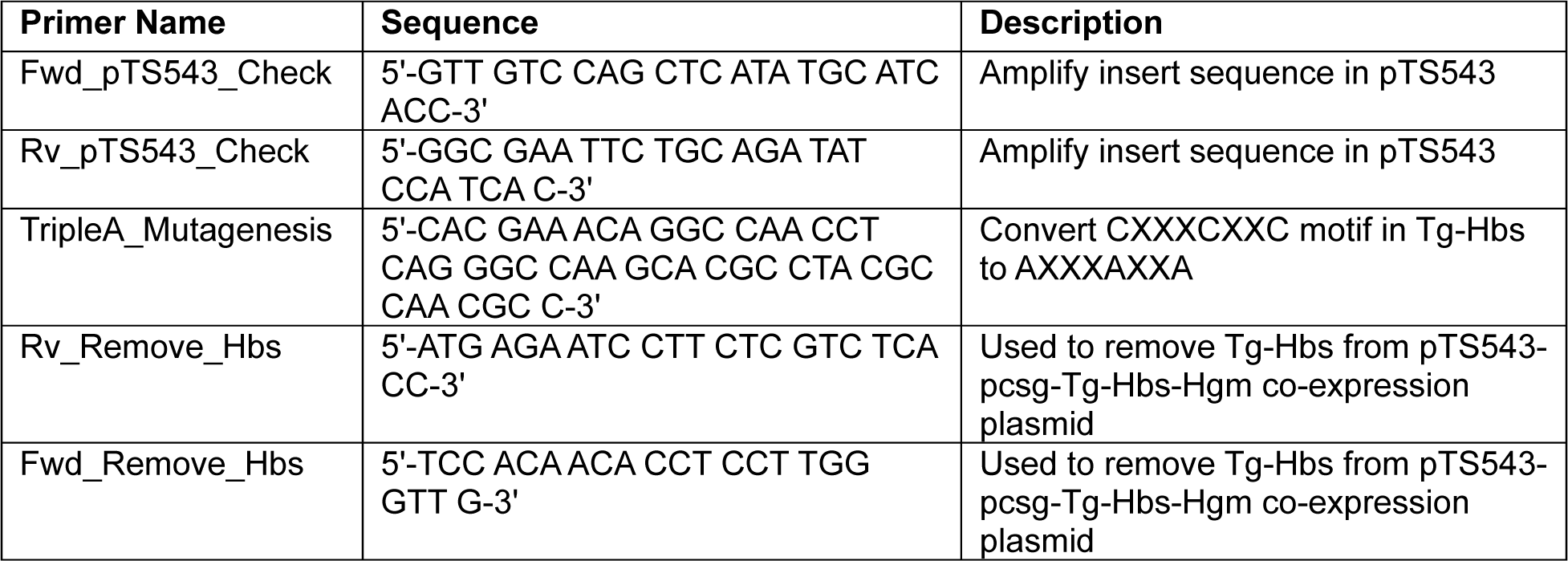
Primers used in this study.

**Figure S1.**
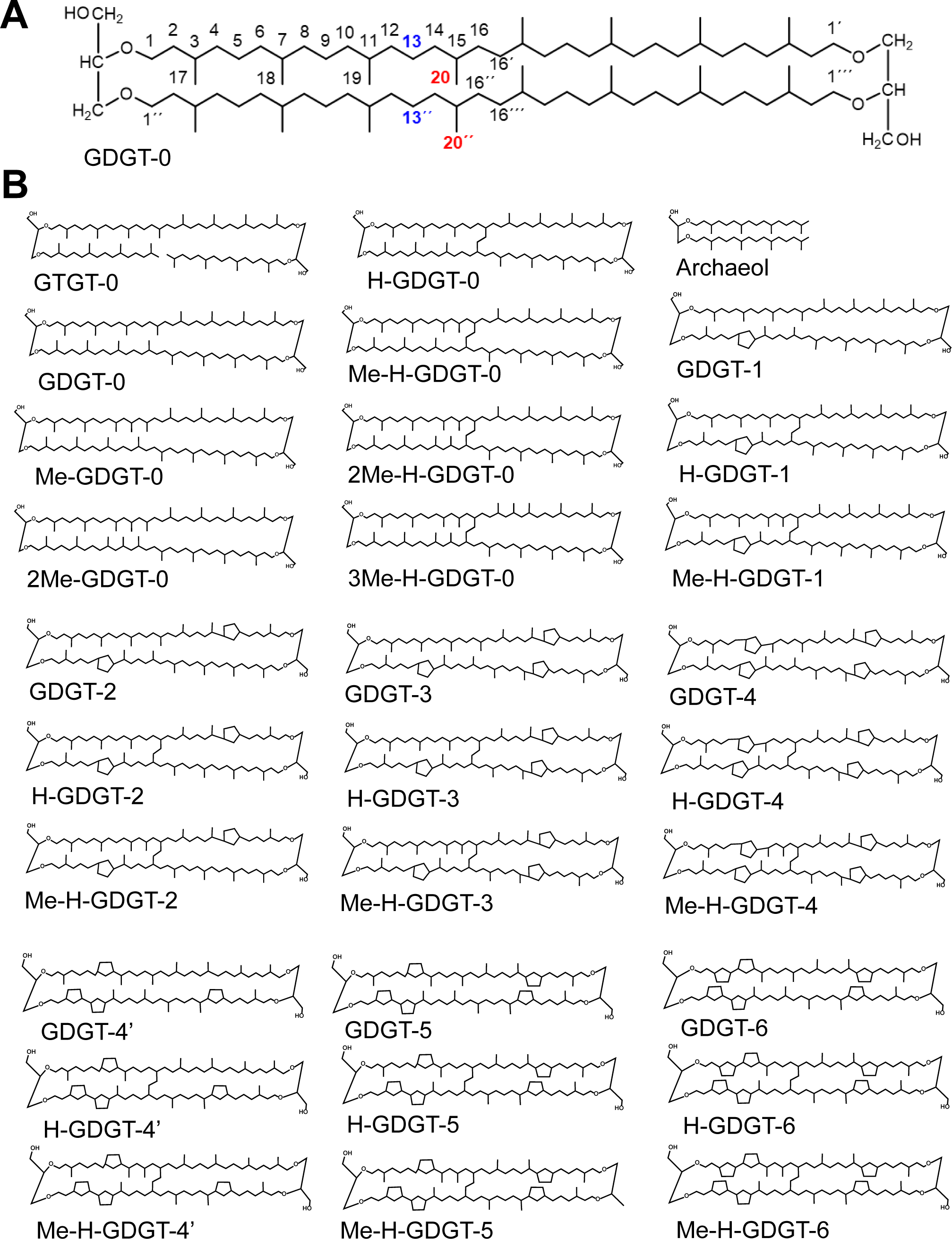
**Archaeal lipids**. A) Numbering scheme of GDGT-0 with the tentative sites of Hbs modification highlighted in red and tentative sites of Hgm modification highlighted in blue. B) Structures of all the lipid compounds analyzed in this study.

**Figure S2.**
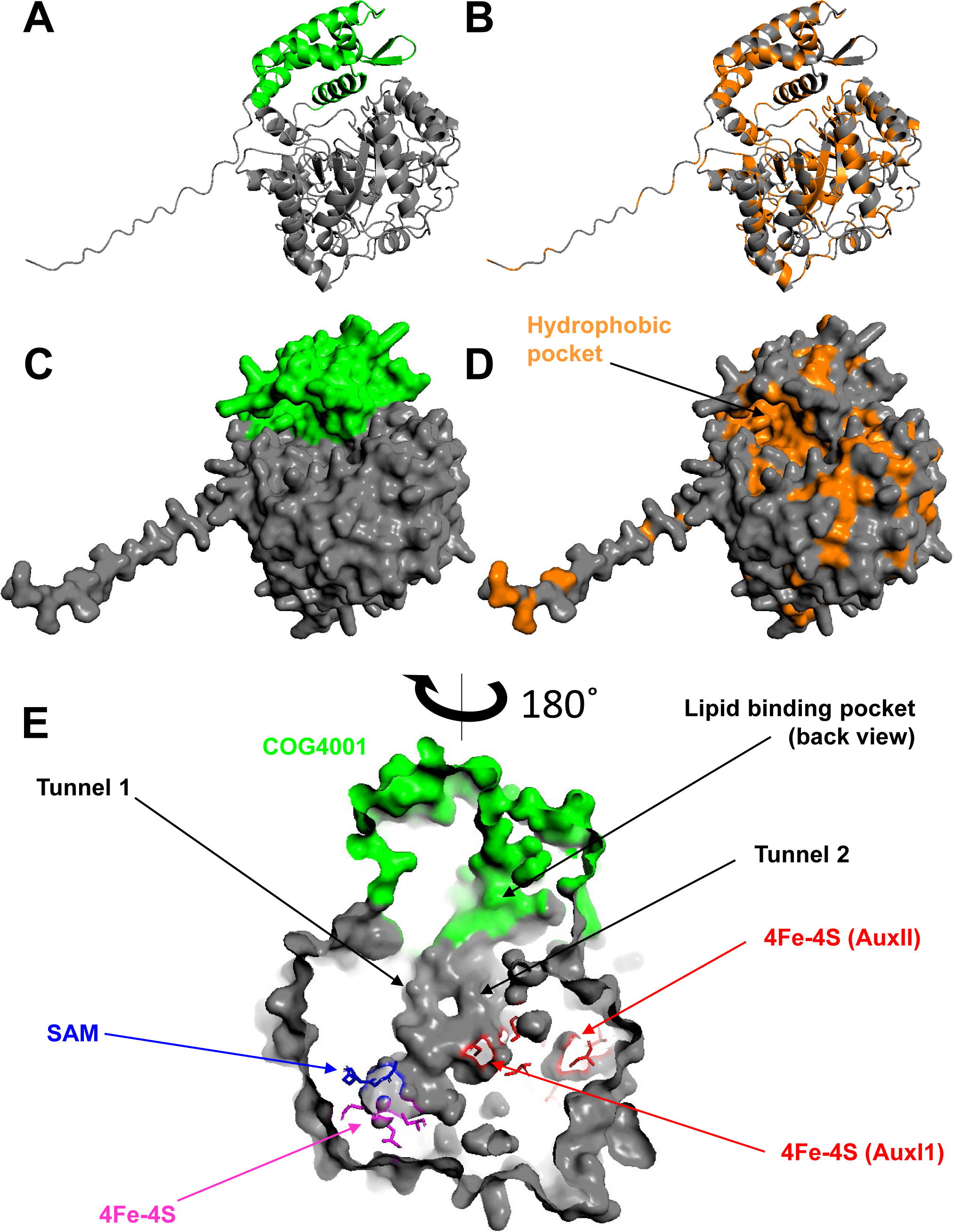
Alpha fold models of Tc-Hbs. A) Alpha fold cartoon structure of Tc-Hbs with COG4001 highlighted in light green. B) Alpha fold cartoon structure of Tc-Hbs with hydrophobic residues highlighted in orange. C) Alpha fold surface structure of Tc-Hbs with COG4001 highlighted in green. Note the pocket formed by COG4001. D) Alpha fold surface structure of Tc-hbs with hydrophobic residues highlighted in orange. Note that the pocket is lined with hydrophobic residues. E) Internal view of Tc-Hbs Alpha fold structure rotated 180 degrees with respect to the above four images. Cysteine residues of the canonical radical SAM 4Fe-4S cluster binding motif (CXXXCXXC) are shown in magenta. SAM binding residues (GGE motif) are shown in blue. Cysteine residues of the SPASM domain responsible for binding the two auxiliary 4Fe-4S clusters (AuxI and AuxII) are shown in red. Note the presence of tunnels connecting the presumed lipid binding pocket to the active site.

**Figure S3.**
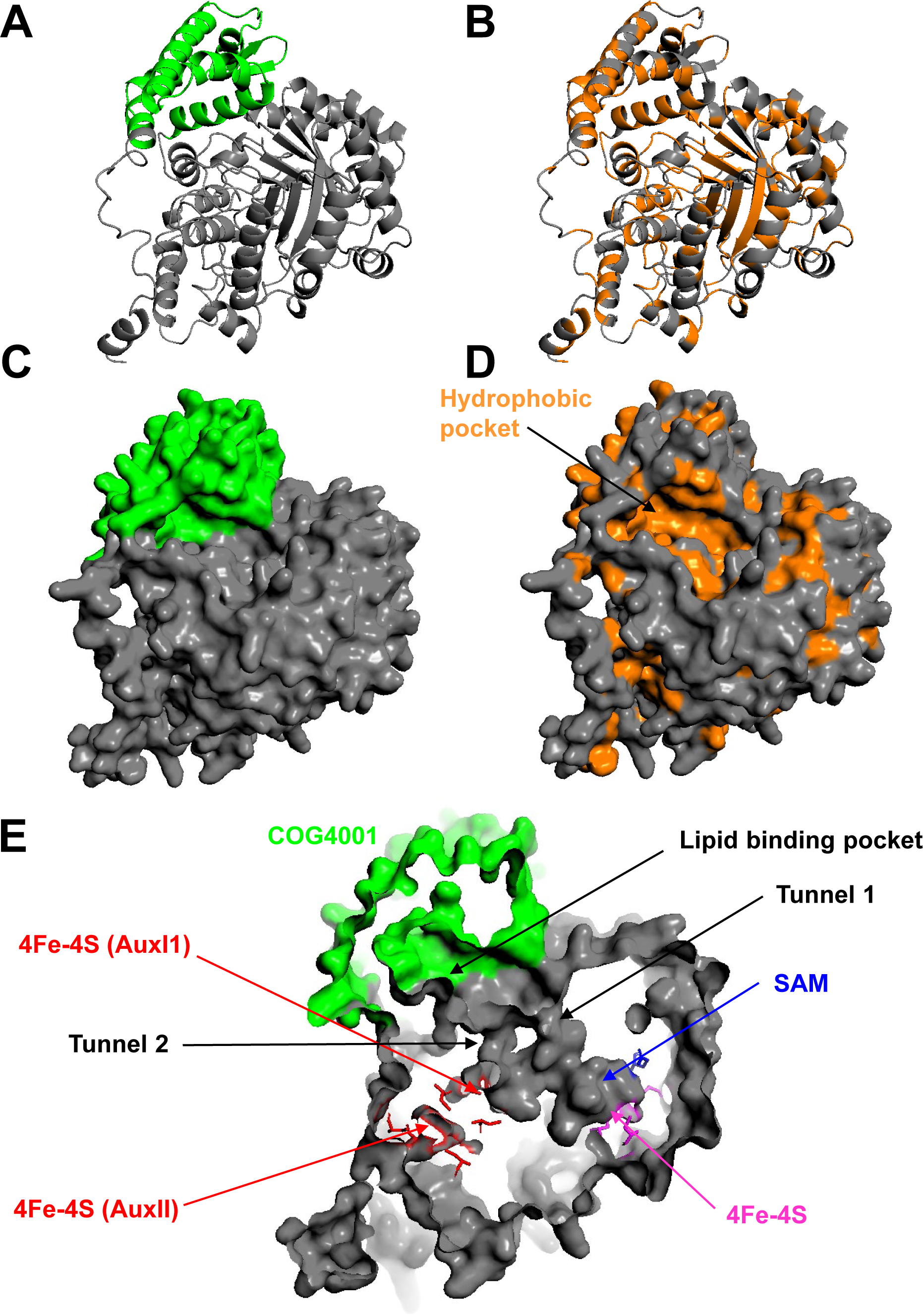
Alpha fold models of Pf-Hbs. A) Alpha fold cartoon structure of Pf-Hbs with COG4001 highlighted in light green. B) Alpha fold cartoon structure of Pf-Hbs with hydrophobic residues highlighted in orange. C) Alpha fold surface structure of Pf-Hbs with COG4001 highlighted in green. Note the pocket formed by COG4001. D) Alpha fold surface structure of Pf-hbs with hydrophobic residues highlighted in orange. Note that the pocket is lined with hydrophobic residues. E) Internal view of Pf-Hbs Alpha fold structure. Cysteine residues of the canonical radical SAM 4Fe-4S cluster binding motif (CXXXCXXC) are shown in magenta. SAM binding residues (GGE motif) are shown in blue. Cysteine residues of the SPASM domain responsible for binding the two auxiliary 4Fe-4S clusters (AuxI and AuxII) are shown in red. Note the presence of tunnels connecting the presumed lipid binding pocket to the active site.

**Figure S4.**
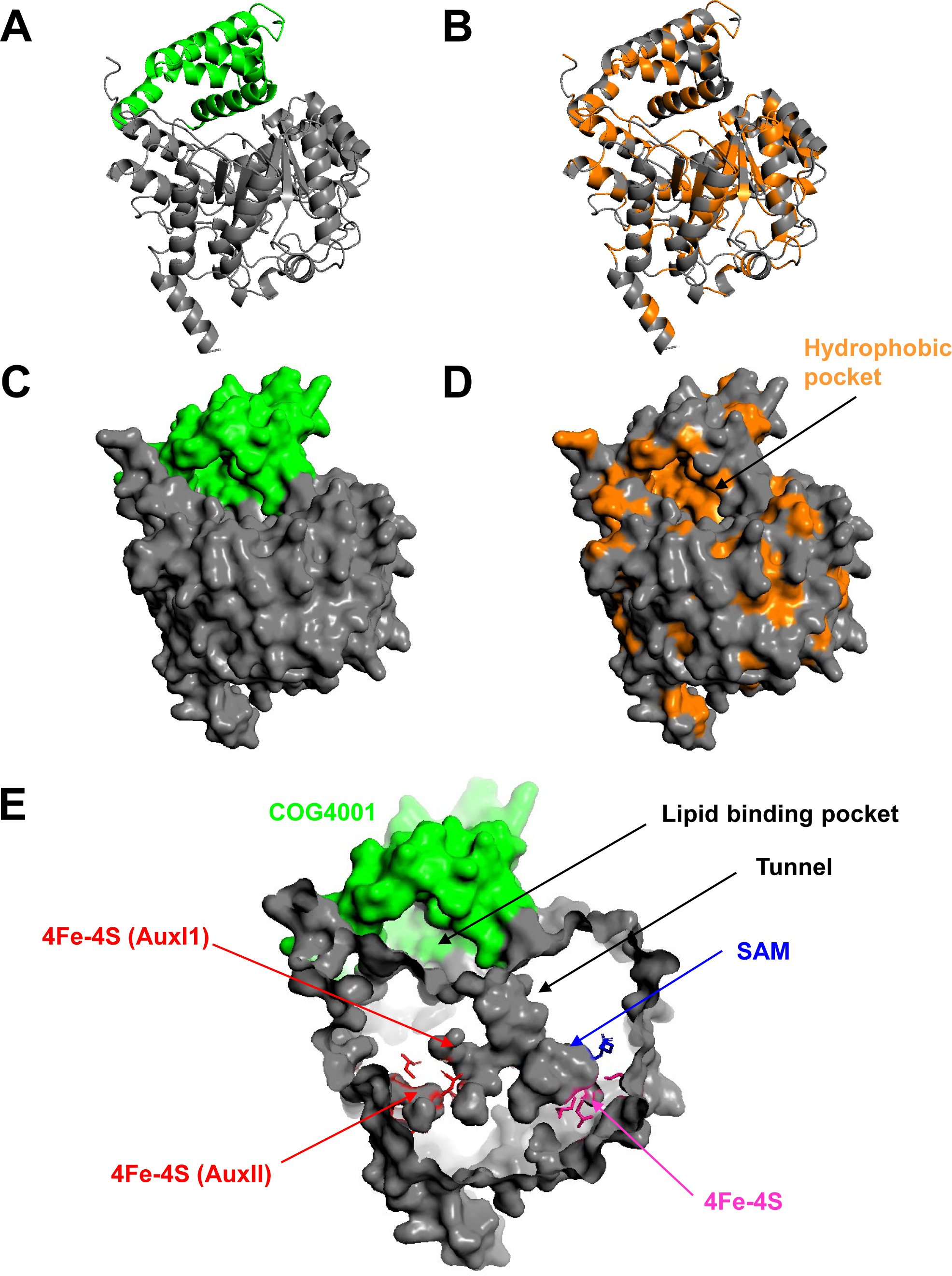
Alpha fold models of Tg-Hbs. A) Alpha fold cartoon structure of Tg-Hbs with COG4001 highlighted in light green. B) Alpha fold cartoon structure of Tg-Hbs with hydrophobic residues highlighted in orange. C) Alpha fold surface structure of Tg-Hbs with COG4001 highlighted in green. Note the pocket formed by COG4001. D) Alpha fold surface structure of Tg-hbs with hydrophobic residues highlighted in orange. Note that the pocket is lined with hydrophobic residues. E) Internal view of Tg-Hbs Alpha fold structure. Cysteine residues of the canonical radical SAM 4Fe-4S cluster binding motif (CXXXCXXC) are shown in magenta. SAM binding residues (GGE motif) are shown in blue. Cysteine residues of the SPASM domain responsible for binding the two auxiliary 4Fe-4S clusters (AuxI and AuxII) are shown in red. Note the presence of a single tunnel connecting the presumed lipid binding pocket to the active site.

**Figure S5.**
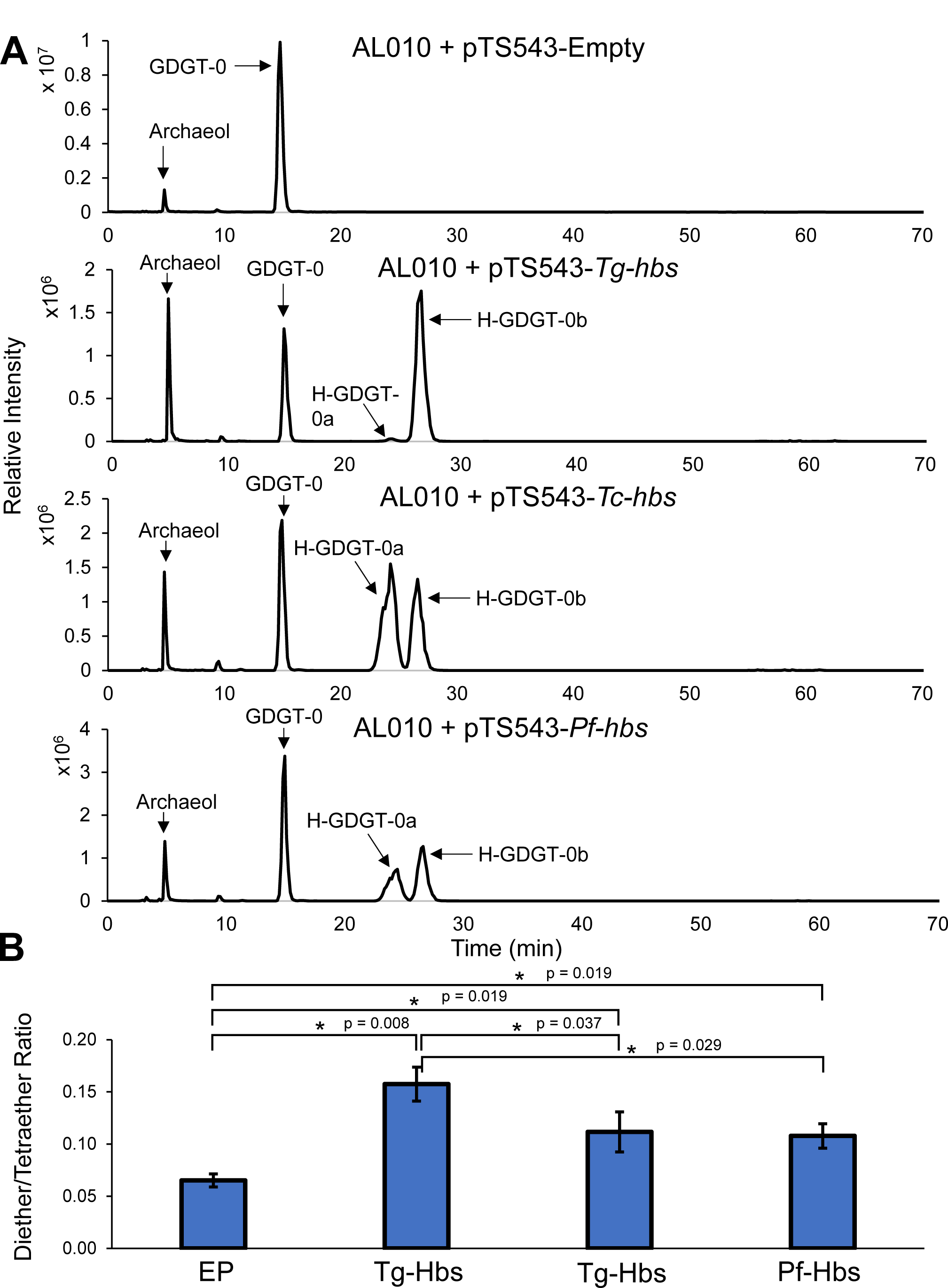
*Thermococcus kodakarensis* increases the relative abundance of diether lipids in response to H-GDGT production. A) Total ion chromatograms (TICs) of the control strain (AL010 + pTS543) and all the *T. kodakarensis* Hbs expression strains analyzed in normal phase. Note the larger archaeol peaks in the Hbs expression strains. B) Bar plot of the diether (bilayer lipid) to tetraether (monolayer lipid) ratio in each strain, showing that the Hbs expression strains increase the ratio of diether to tetraether lipids relative to the control and that the Tg-Hbs expression strain increases them more than Tc-Hbs and Pf-Hbs expression strains. * Denotes significant differences and associated p-values are shown adjacently.

**Figure S6.**
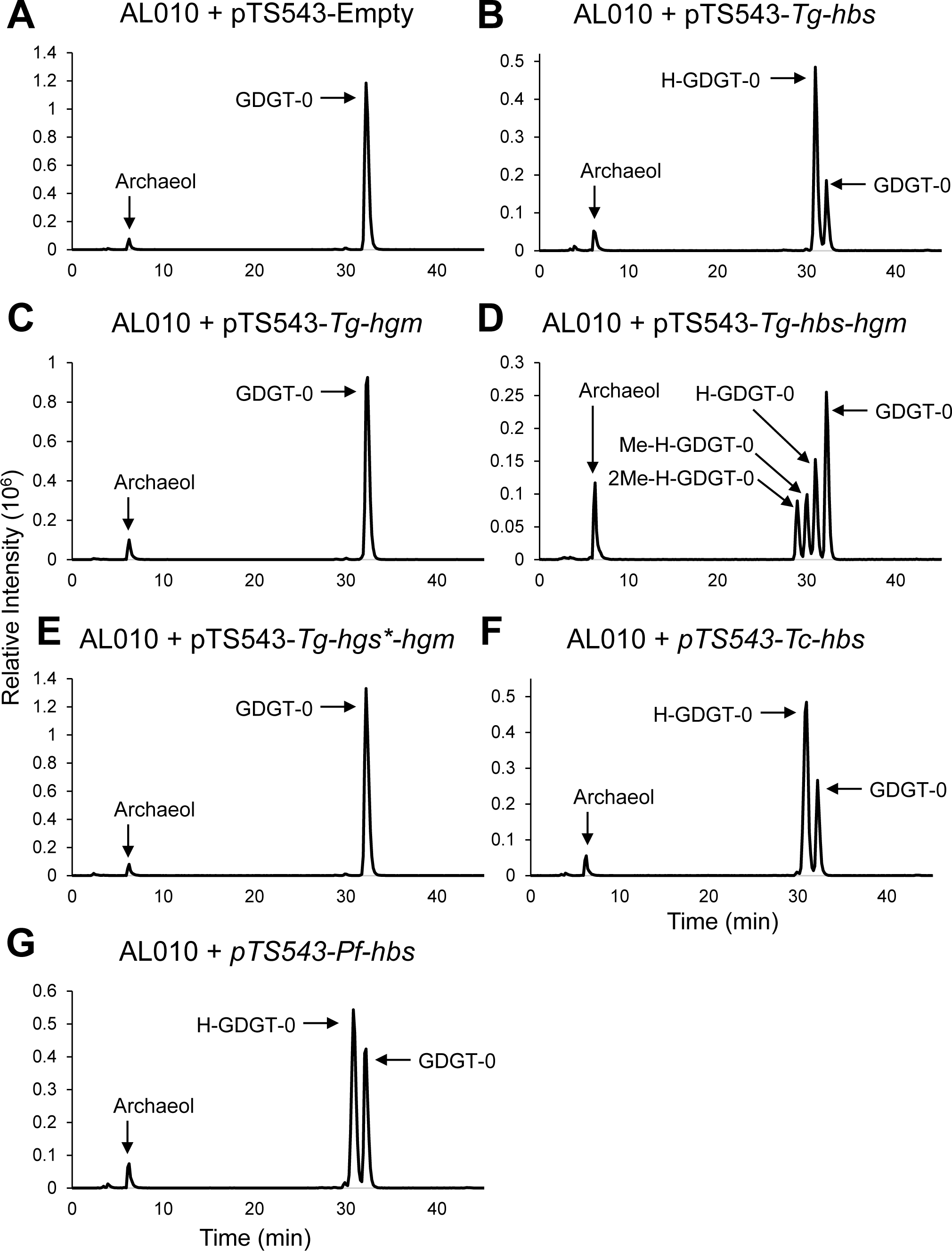
Total Ion Chromatograms (TICs) of all *T. kodakarensis* strains examined in reverse phase. A) TIC of the control strain showing the presence of only GDGT-0 and archaeol. B) TIC of Tg-Hbs expression strain showing the presence of H-GDGT-0, GDGT- 0, and archaeol. C) TIC of Tg-Hgm expression strain showing the presence of GDGT-0 and archaeol. Me-GDGT-0 and 2Me-GDGT-0 peaks are too small to see and overlap with the GDGT-0 peak. D) TIC of the Tg-Hbs + Tg-Hgm co-expression strain showing the presence of 2Me-H-GDGT-0, Me-H-GDGT-0, H-GDGT-0, GDGT-0, and archaeol. 3Me- H-GDGT-0 co-elutes with 2Me-H-GDGT-0 and cannot be seen. E) TIC of catalytically inactivated Tg-Hbs* + Tg-Hgm co-expression strain showing the presence of GDGT-0 and archaeol and the absence of H-GDGT-0, indicating Hbs was successfully inactivated. Me-GDGT-0 and 2Me-GDGT-0 peaks are too small to see and overlap with the GDGT-0 peak. F) and G) TICs of Tc-Hbs and Pf-Hbs expression strains, respectively, showing presence of H-GDGT-0, GDGT-0, and archaeal. These two TICs also show that the H-GDGT-0a and H-GDGT-0b isomers (which are present in near 1:1 amounts in Tc-Hbs and Pf-Hbs expression strains) co-elute during reverse phase chromatography.

**Figure S7.**
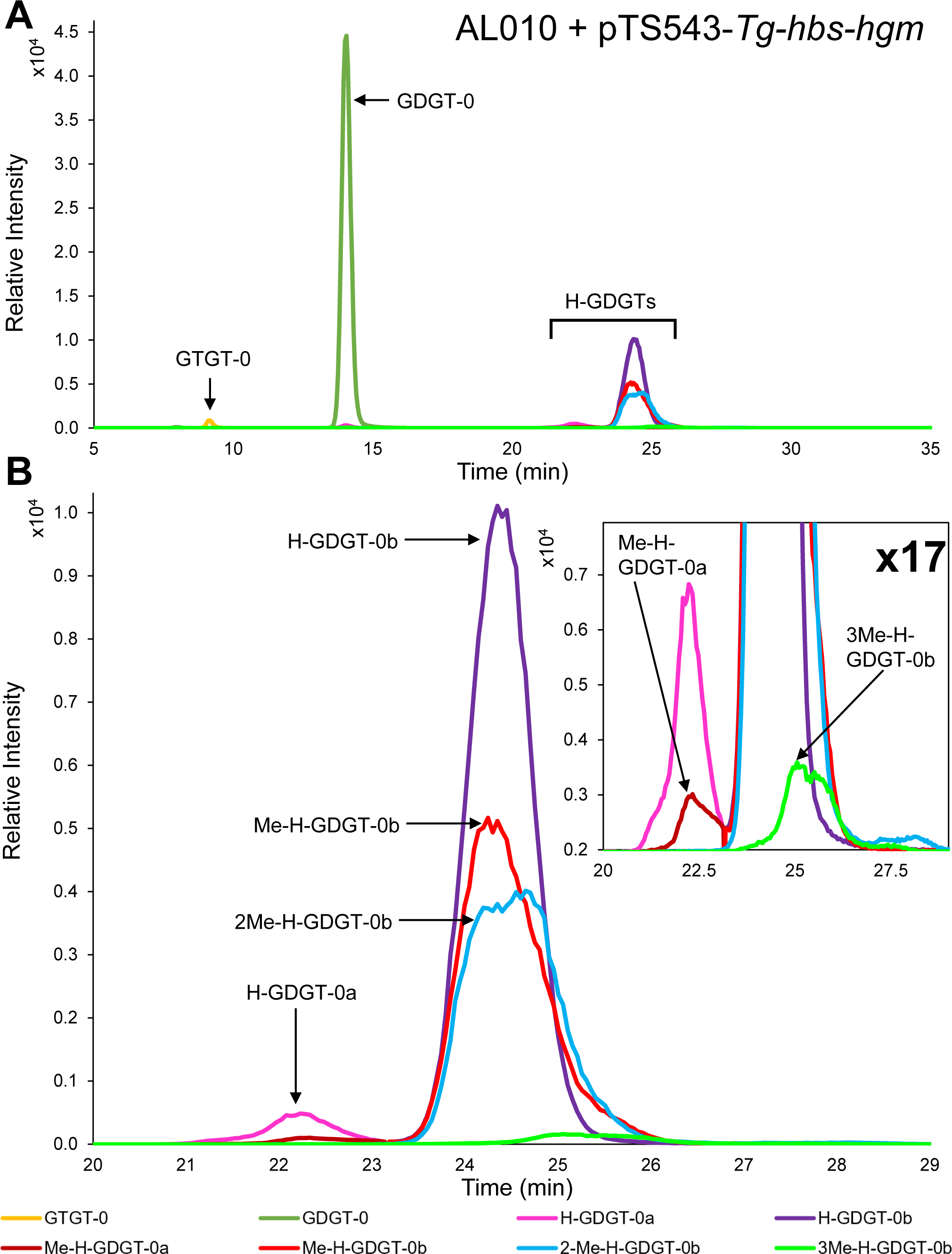
Methylated H-GDGTs co-elute with their unmethylated counterparts during normal phase chromatography. A) Merged extracted ion chromatograms (EIC) of the Tg-Hbs + Tg-Hgm overexpression strain, showing the presence of GTGT-0, GDGT- 0, and a variety of methylated and unmethylated H-GDGTs that co-elute. B) Close up of the H-GDGTs seen in A. Both the H-GDGT-0a and H-GDGT-0b isomers have co-eluting peaks, corresponding to the mono-methylated version of each isomer. However, di- methylated and tri-methylated peaks only occur under the more abundant H-GDGT-0b isomer, revealing an absence of di- and tri-methylation of H-GDGT-0b. The absence of di-methylation on H-GDGT-0a is intriguing as di-methylated GDGTs are observed during Tg-Hgm alone expression. In both of these cases, the mono-methylated substrate is present in low abundance (small amounts of Me-H-GDGT-0a and small amounts of Me- GDGT-0). However, a second methylation occurs on Me-GDGT-0 but not Me-H-GDGT- 0a, potentially suggesting the location of the bridge in the H-GDGT-0a isomer blocks the addition of a second methyl group by Hgm.

**Figure S8.**
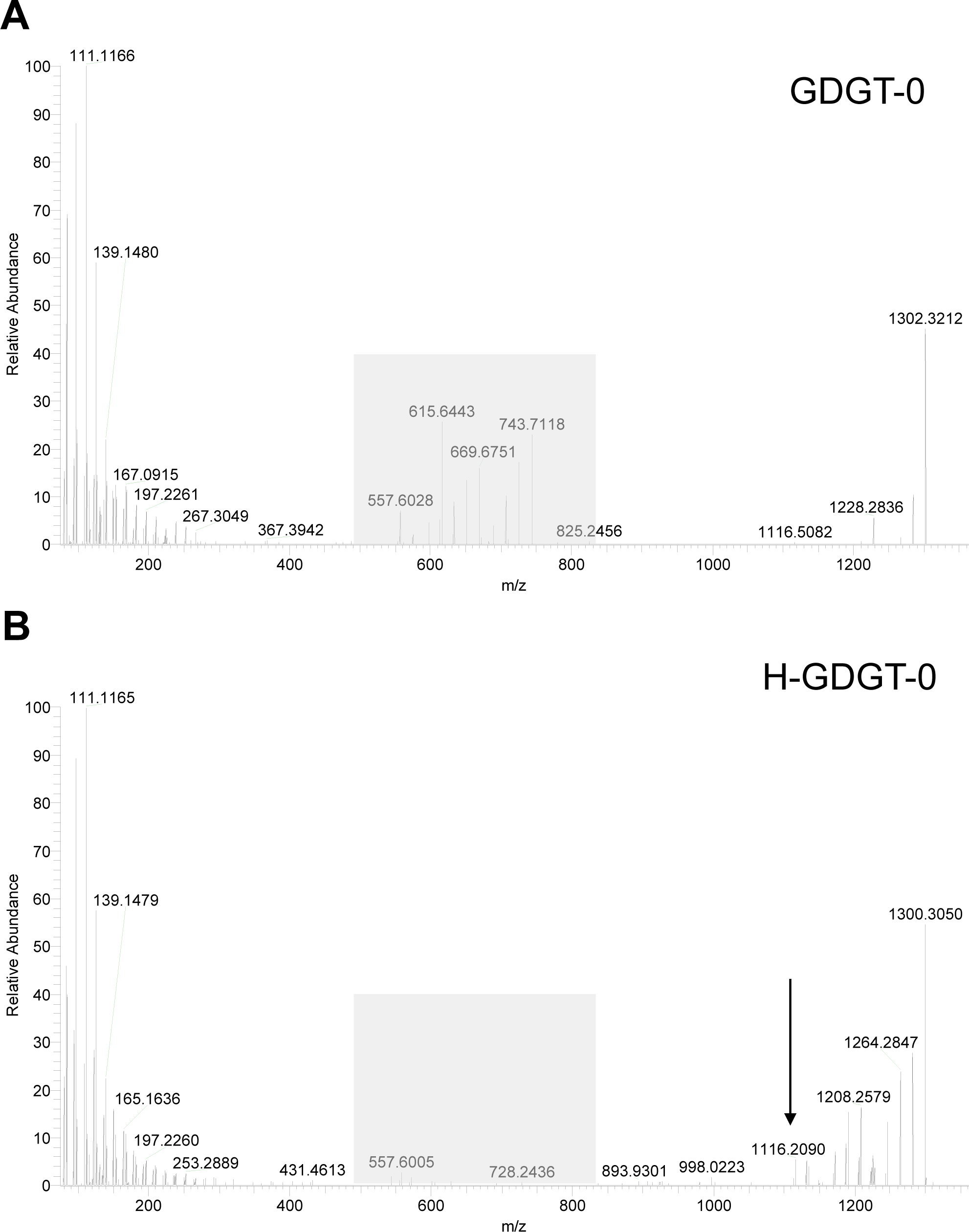
Validating the identity of H-GDGT-0 with LC-MS/MS. A) MS^2^ spectra of GDGT-0. Note the presence of fragments corresponding to the loss of a single biphytanyl tail in the grey box. B) MS^2^ spectra of reverse phase co-eluted H-GDGT-0a and H-GDGT- 0b isomers from the Tg-Hbs expression strain. Fragments in the gray box are diagnostically absent in the H-GDGT-0 spectra due to the cross-linking of the biphytanyl chains to form a monoalkyl chain, preventing loss of single biphytanyl tails. The arrow indicates the diagnostic product ion (m/z = 1116.2) for H-GDGT-0, corresponding to the free monoalkyl chain.

**Figure S9.**
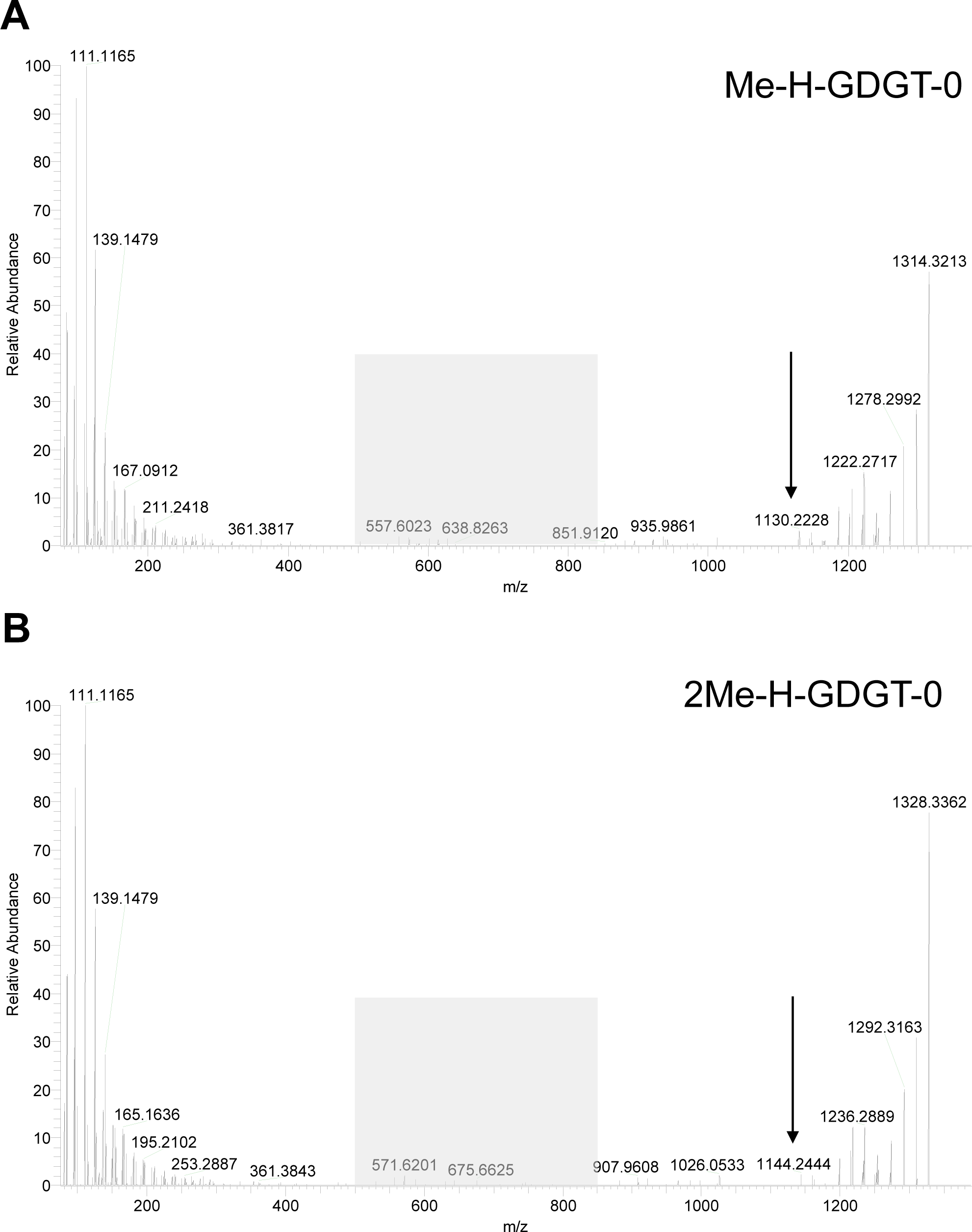
Confirmation of methylation within the monoalkyl core of H-GDGTs with LC-MS/MS. A) MS^2^ spectra of Me-H-GDGT-0. Fragments corresponding to the loss of a single biphytanyl tail in the gray box are diagnostically absent, indicating this compound is an H-GDGT. The arrow indicates the diagnostic product ion (m/z = 1130.2) for Me-H-GDGT-0, corresponding to the free monoalkyl chain with an additional methylation as indicated by the +14-mass shift in this product ion compared to unmethylated H-GDGT-0. Thus, Hgm introduces methylations on the lipid tails rather than the glycerol backbone as is seen in some archaea. B) MS^2^ spectra of 2Me-H-GDGT-0. Fragments corresponding to the loss of a single biphytanyl tail in the gray box are diagnostically absent, indicating this compound is an H-GDGT. The arrow indicates the diagnostic product ion (m/z = 1144.2) for 2Me-H-GDGT-0, corresponding to the free monoalkyl chain with two additional methylations as indicated by the +28 mass shift in this product ion compared to unmethylated H-GDGT-0. Note that a +28 mass shift could also indicate single ethylation (-CH_2_CH_3_) rather the double methylation. The spectra of 2Me-GDGT-0 in Figure S10 supports the notion of double methylation rather than single ethylation.

**Figure S10.**
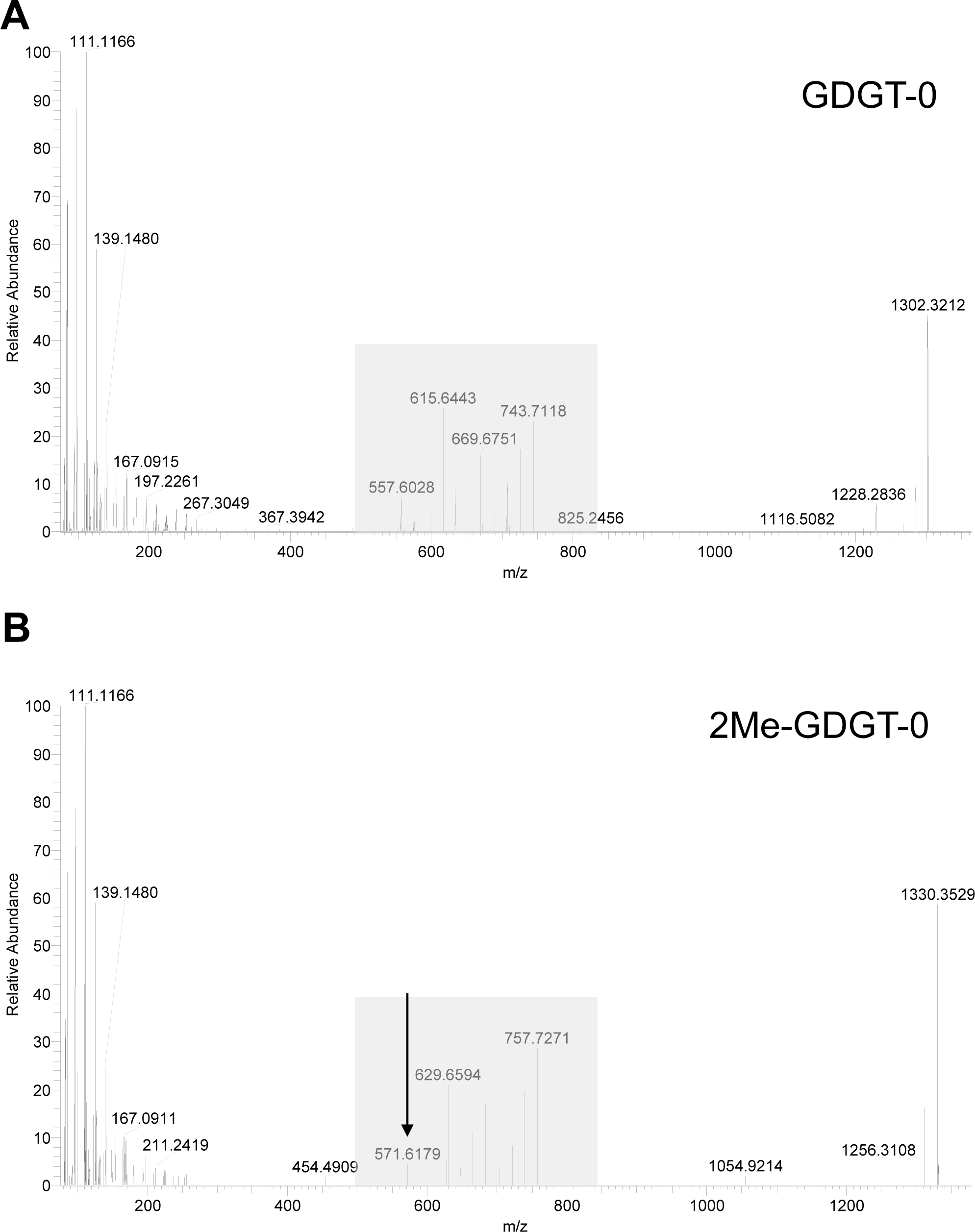
2Me-GDGT-0 possesses two methylations, one on each opposing biphytanyl tail. A) MS^2^ spectra of GDGT-0. Note the presence of fragments corresponding to the loss of a single biphytanyl tail in the grey box. B) MS^2^ spectra of 2Me-GDGT-0 from the Tg-Hgm expression strain. Fragments in the gray box are present indicating this compound is a GDGT. The arrow indicates the diagnostic product ion (m/z = 571.6) for 2Me-GDGT-0, corresponding to a single biphytanyl chain which has been cleaved off from the rest of the molecule. This mass of this biphytanyl tail is shifted by +14 compared to the biphytanyl chain released in GDGT-0 (m/z =557.6), demonstrating the methylation occurs in the hydrocarbon chain and not the glycerol backbone. Further, no peaks corresponding to the release of a biphytanyl tail with two methylations (or a single ethylation) on it are detected (no m/z = 585.6 peaks), indicating that Hgm adds its second methyl group to the tail opposite of where the first was added.

**Figure S11.**
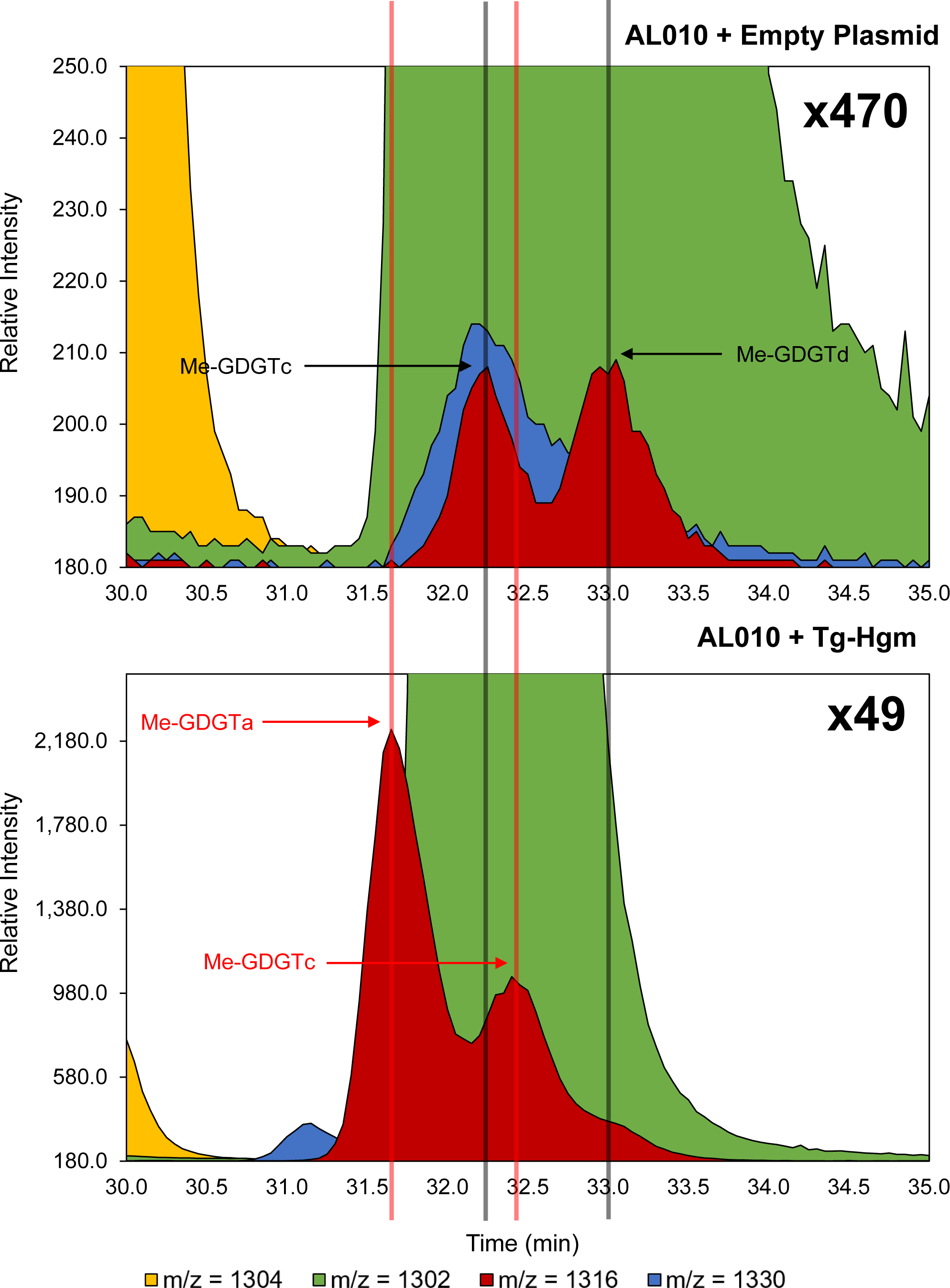
Mono-methylated GDGTs are present but differ in the control and Tg- Hgm expression strains. Merged EICs showing the different chromatographic behavior of mono-methylated GDGTs in the control strain (AL010 + Empty Plasmid (EP)) and the Tg-Hgm expression strain. Both strains produce Me-GDGTs: two isomers termed Me- GDGTb and Me-GDGTd in the control strain and, two different isomers termed Me- GDGTa and Me-GDGTc in the Hgm expression strain. The elution times of all isomers differ as indicated by the lines delineating their peak apex points. Note that the control strain is shown at 470x the height of the GDGT-0 (m/z = 1302) peak while Hgm expression strain is shown at 49x the height of the GDGT-0 (m/z = 1302) peak, demonstrating more robust GDGT methylation in the Hgm expression strain by an order of magnitude. The structural differences between the isomers are not known.

**Figure S12.**
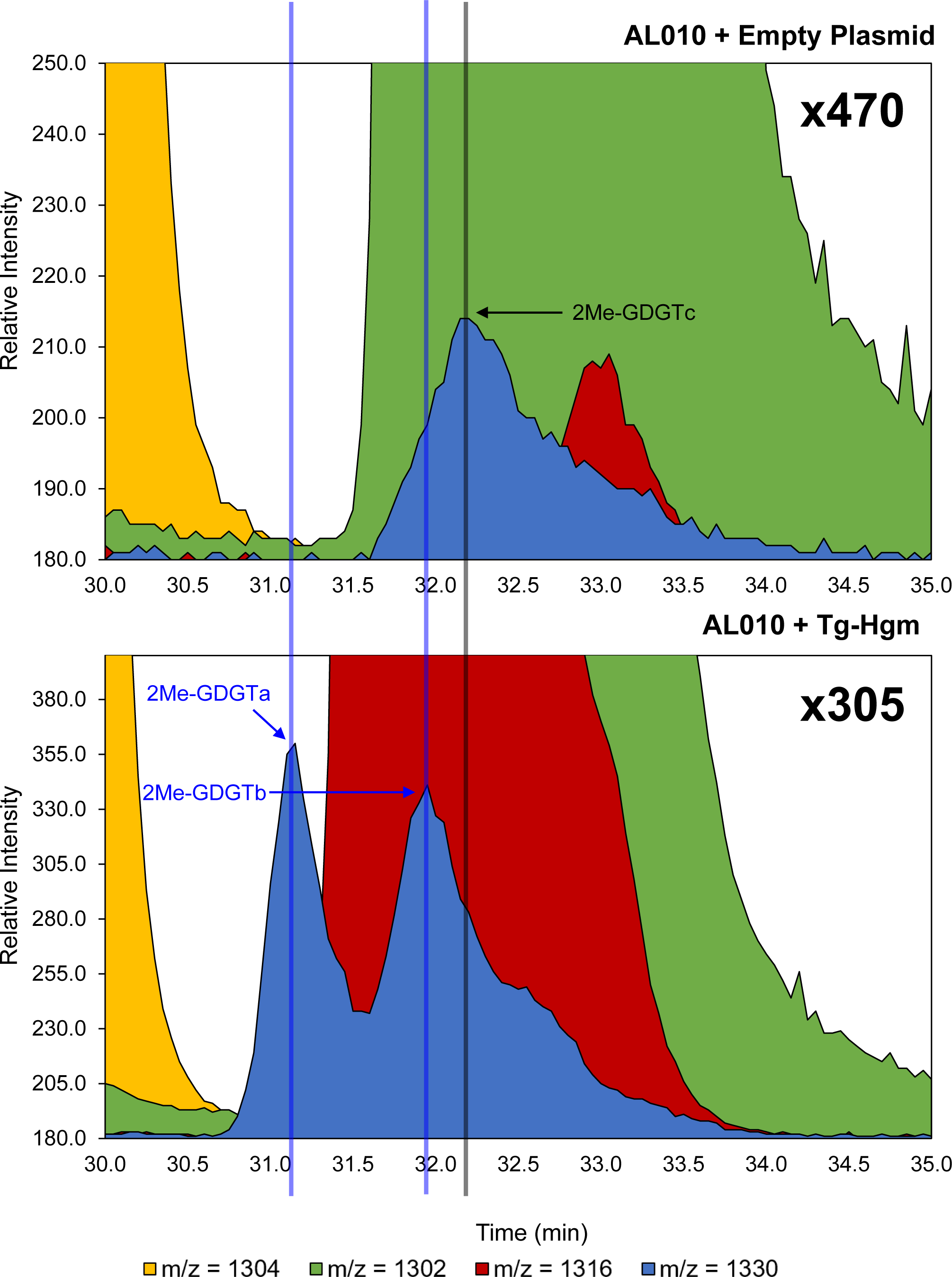
Di-methylated GDGTs are present but differ in the control and Tg-Hgm expression strains. Merged EICs showing the different chromatographic behavior of di- methylated GDGTs in the control strain (AL010 + Empty Plasmid (EP)) and the Tg-Hgm expression strain. The control strain produces one di-methylated GDGT isomer (2Me- GDGTc) while the Hgm expression strain produces two different 2Me-GDGT isomers (2- Me-GDGTa and 2Me-GDGTb). The elution times of all isomers differ as indicated by the lines delineating the peak apex points. The control strain is shown at 470x the height of the GDGT-0 (m/z = 1302) peak. The Hgm expression strain is shown at 305x the height of the GDGT-0 (m/z = 1302) peak. The structural differences between the isomers are not known.

**Figure S13.**
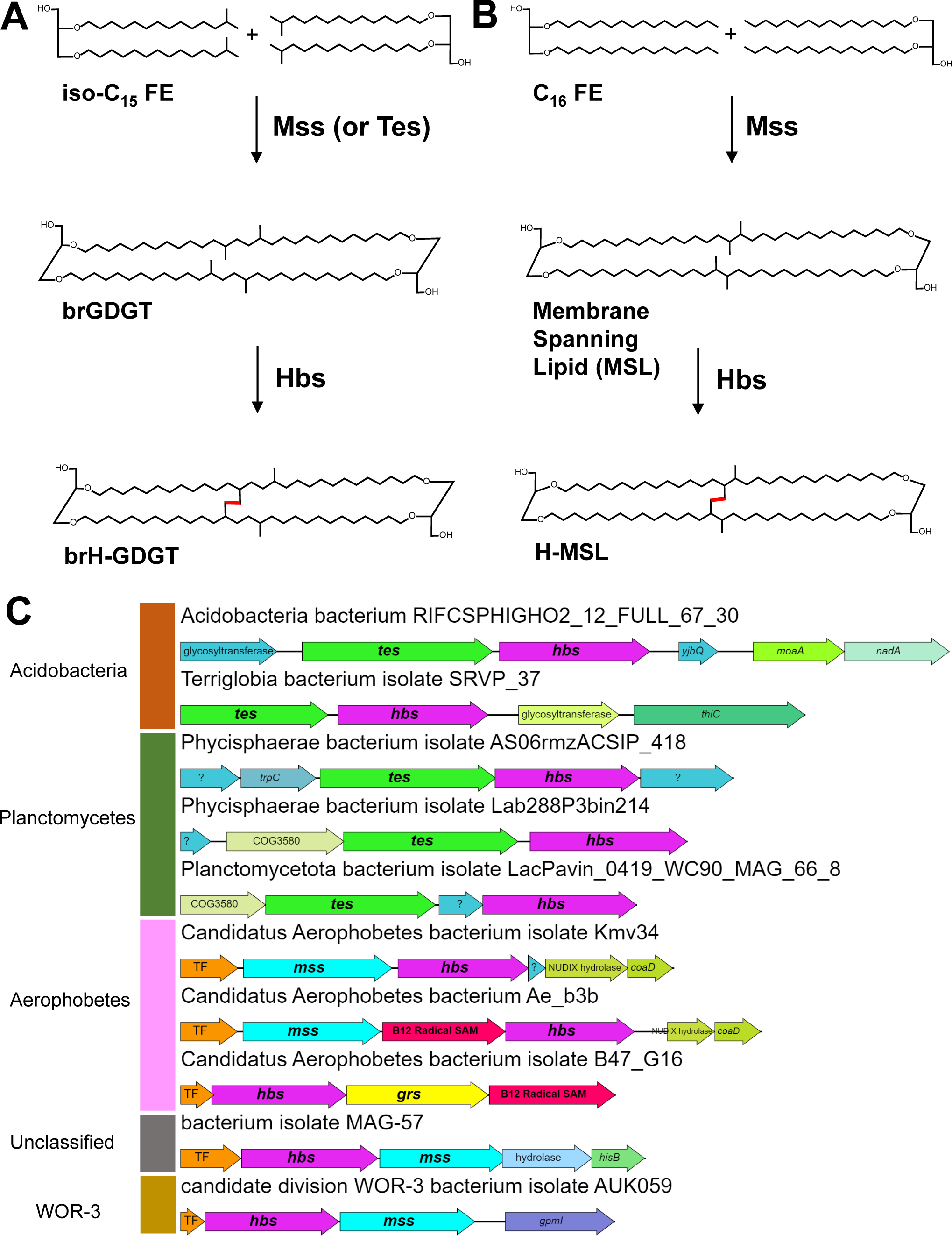
Bacterial Hbs homologs may be responsible for the synthesis of brH- GDGTs and other bacterial H-shaped membrane spanning lipids. A) Hypothesized biosynthetic pathway for the formation of brH-GDGTs. Mss or Tes may dimerize two iso- C_15_ fatty ethers (FE) to form a brGDGT. Hbs may then bridge the two alkyl chains together at their branching points to form a brH-GDGT. B) Hypothesized biosynthetic pathway for the formation of H-shaped membrane spanning lipids (H-MSLs) not based on iso-diabolic acid. Mss may dimerize two C_16_ fatty ethers to form a membrane spanning lipid like those seen in *Thermotoga maritima*. Hbs may then bridge the two alkyl chains together at their branch points to form a H-MSL. C) Bacterial *hbs* genes occur in gene clusters with *mss* and *tes* in diverse bacterial phyla including Acidobacteria, Planctomyectes, Aerophobetes, WOR-3, and unclassified bacteria. Hbs primarily clusters with tes in Acidobacteria and Planctomycetes while it primarily clusters with *mss* in candidate bacterial phyla such as the Aerophobetes. Note the interesting presence of *grs* homologs in the Aerophobetes gene clusters as well, suggesting they may be capable of producing cyclized brH-GDGTs

**Figure S14.**
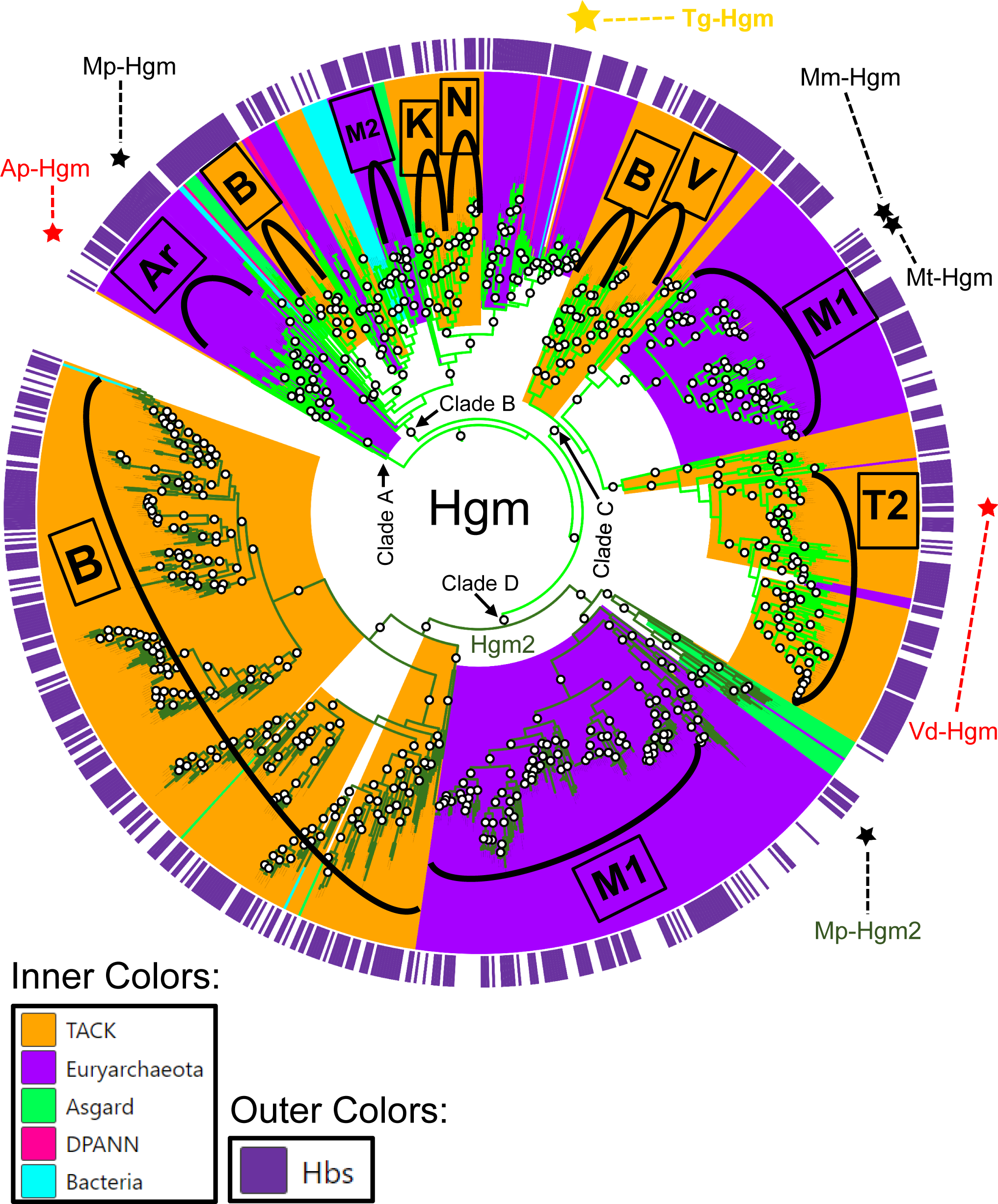
Hgm homologs are found in diverse archaea dominated by the Bathyarchaeota, Methanomada, and Thermoprotei. Hgm tree showing the phylogenetic distribution of Hgm homologs and their co-occurrence with Hbs genes. Inner ring colors denotate the major phyla each Hgm sequence belongs to. More specific phylogenetic clades are denoted as follows: Archaeoglobi [Ar], Bathyarchaeota [B], Korarchaeota [K], Methanomada [M1], Methanomicrobia [M2], Nitrososphaeota [N], Thermoproteota [T2], and Verstraetearchaeota [V]. The Hgm protein expressed in this study is denoted by a gold star. The Hgm proteins belonging to archaea we show produce methylated H-GDGTs in this work are denoted by red stars: *Archaeoglobus profundus* Hgm (Ap-Hgm) and *Vulcanisaeta distributa* Hgm (Vd-Hgm). Other archaea which have been demonstrated to produce methylated H-GDGTs in culture are denoted by the black stars: *Methanopyrus kandleri* Hgm (Mp-Hgm), *Methanothermobacter marburgensis* Hgm (Mm-Hgm), and *Methanothermobacter thermautotrophicus* Hgm (Mt-Hgm). The outer purple ring corresponds to the presence of an Hbs gene within the genomes of the Hbs encoding microbes. Clades A, B, and C that are shown with light green branch colors correspond to Hgm 1 while clade D, shown with dark green branch colors, corresponds to Hgm2. Note that the Hgm2 clade is almost exclusively composed of sequences from Bathyarchaeota and Methanomada.

**Figure S15.**
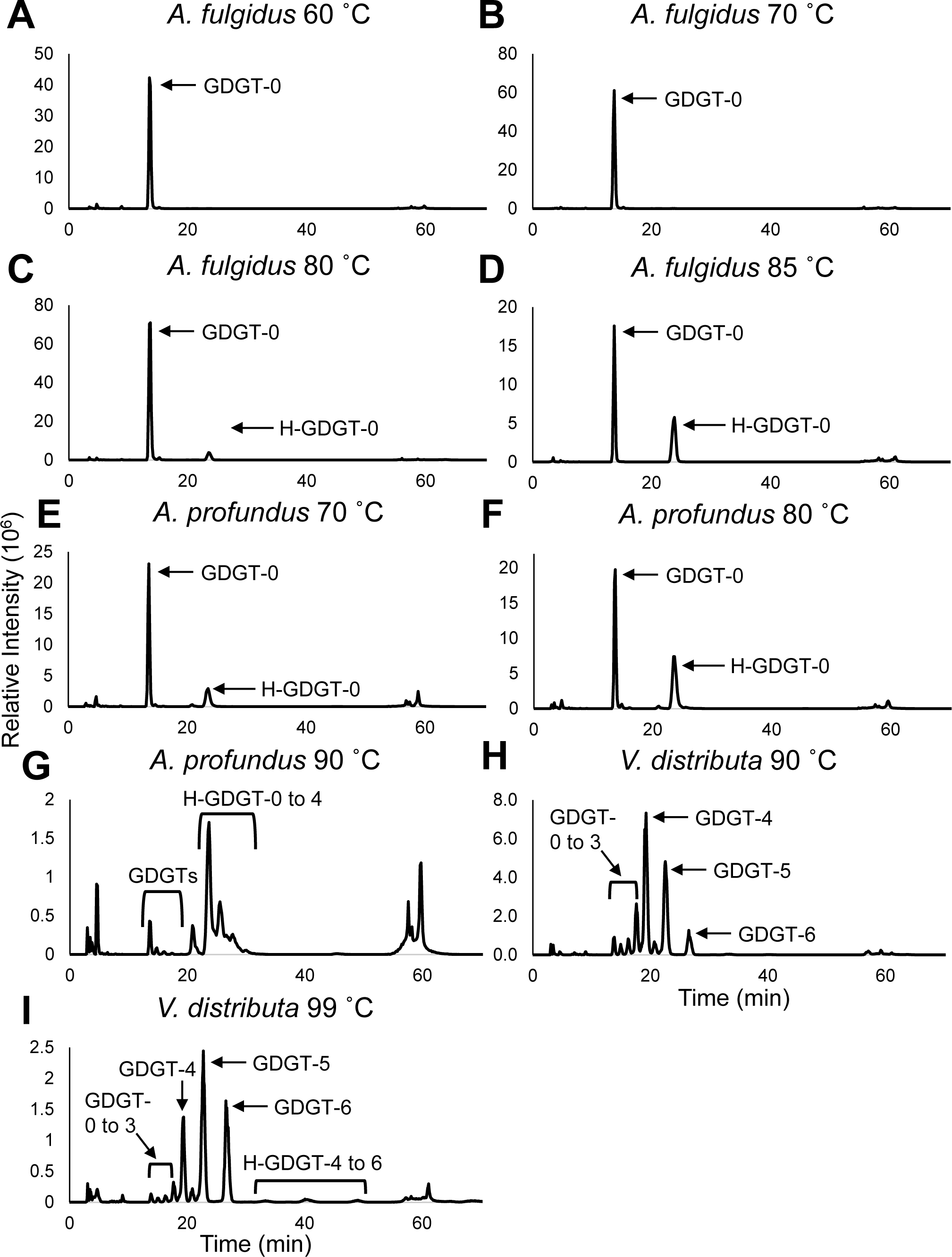
Total ion chromatograms (TICs) of *A. fulgidus*, *A. profundus*, and *V. distributa* lipids across all temperatures analyzed. TICs of *A. fulgidus* grown at A) 60°C, B) 70°C, C) 80°C, and D) 85°C, respectively, showing the presence of GDGT-0 and the appearance of H-GDGT-0 at 80°C and 85°C. TICs of *A. profundus* grown at E) 70°C, F) 80°C, and G) 90°C, respectively, showing the presence of diverse monolayer lipid species and the shift from GDGT dominance at 70 °C and 80 °C to H-GDGT dominance at 90 °C. TICs of *V. distributa* grown at H) 90 °C and I) 99 °C, respectively, showing the presence of multiple cyclized GDGTs. H-GDGTs with 4, 5, and 6 rings are marginally visible in panel I.

**Figure S16.**
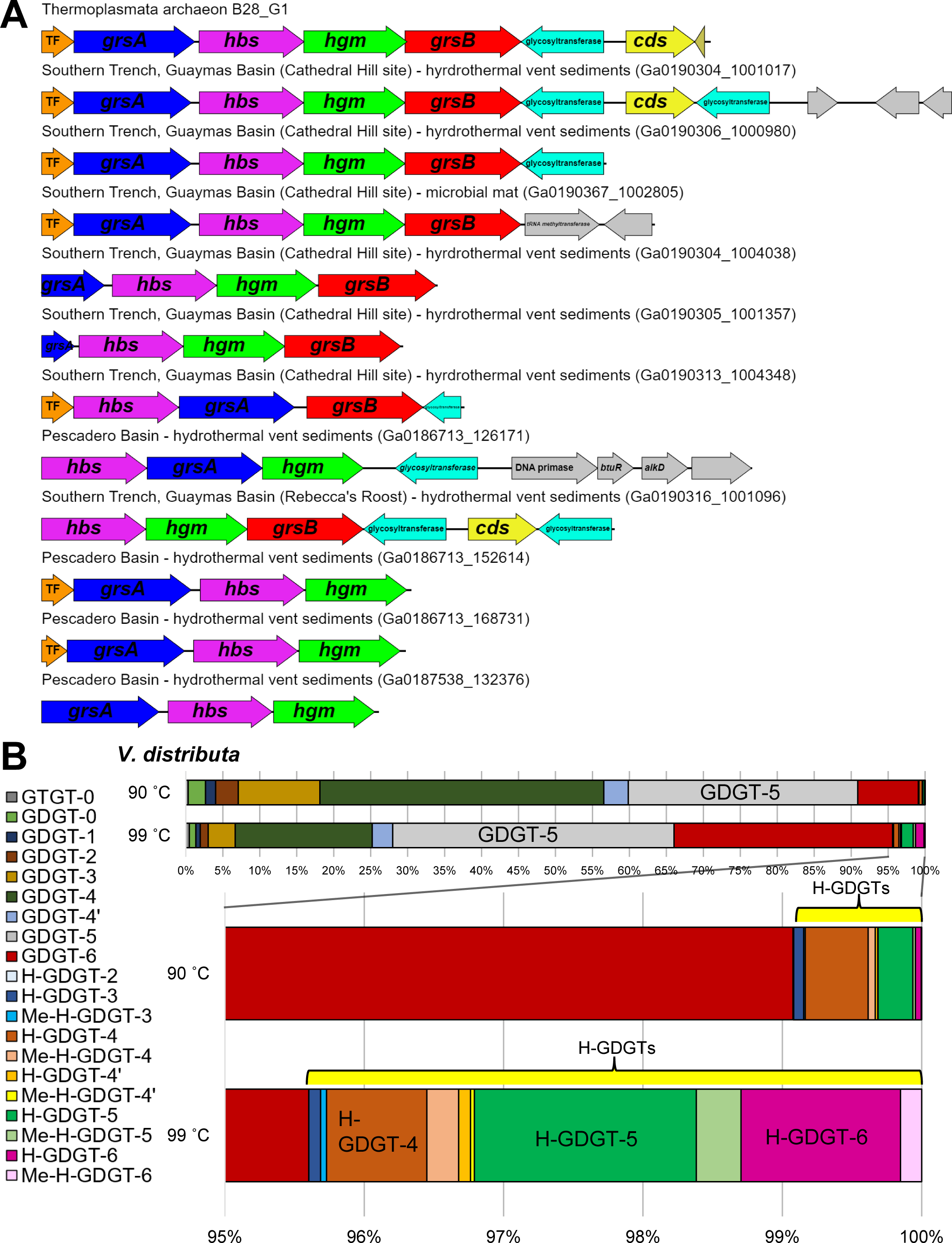
Highly cyclized H-GDGT production in archaea. A) *Thermoplasmata archaeon* B28_G1 related scaffolds from Guaymas and Pescadero Basin showing the clustering of *grsA*, *hbs*, *hgm*, and *grsB* genes, suggesting the presence of operons dedicated to highly cyclized H-GDGT production and methylation. Note the presence of calditol synthase (Cds) as well on some of these scaffolds. Cds is involved in the biosynthesis of an archaeal lipid headgroup which protects cells against acidic pH (4) – its presence here may indicate that these highly cyclized GDGTs are synthesized in response to exposure to acidic pH. TF = transcription factor. B) Relative of abundance of the diverse monolayer lipids found in *V. distributa*.

